# Disinhibition of a recurrent attractor gates a persistent goal signal for navigation

**DOI:** 10.1101/2025.10.07.681003

**Authors:** Aaron J. Lanz, Nicholas D. Kathman, Emily Hao, Bard Ermentrout, Katherine I. Nagel

## Abstract

Recurrent attractor networks are widely thought to form the basis of working memory^1–3^, but how stable attractor activity can be rapidly switched on and off is unclear^4–7^. Here we investigate how stability and rapid switching can be combined in a discrete recurrent circuit of the fly navigation center^8^. hΔK and PFG neurons are recurrently connected in a ring structure. Using *in vivo* imaging, we find that these two populations exhibit shared persistent bump activity that turns on with odor and terminates at the end of a goal-directed upwind run. Using whole-cell recordings, we show that persistence in hΔK depends on recurrent signalling, and that hΔK receives slow recurrent excitation and fast inhibition from its synaptic partners. Computational modeling reveals that this combination of slow excitation with fast inhibition yields persistent attractor dynamics over a range of excitation and inhibition strengths. Next we examine the mechanisms that allow this activity bump to be rapidly turned on and off. We find that while both populations show positionally stable bump activity during goal-directed runs, during turns and rest the PFG bump tracks heading while hΔK is supressed. We can reproduce these differential dynamics in our model by using inhibition to dynamically uncouple activity in hΔK from PFG. When hΔK is inhibited, PFG neurons follow their inputs from the compass system; when hΔK is disinhibited, recurrent interactions lock this input into place, forming a heading memory. Consistent with this model, we find that inhibitory inputs onto hΔK increase during turns and are suppressed during odor input and goal-directed upwind runs. Our work reveals how disinhibition can serve as a gate to rapidly write an ongoing measurement to a recurrent memory circuit.

## Introduction

Distributed networks of recurrently connected neurons are widely thought to form the substrate for short-term working memory^1–3^. During working memory tasks, neurons in the prefrontal cortex and elsewhere exhibit persistent activity that turns on with cue presentation and terminates with behavior^9–10^. The duration of persistence in these neurons is much longer than their intrinsic membrane time constants, arguing that persistence arises from network interactions^3^. Computational models that feature local recurrent excitation and global inhibition can recreate the types of persistent activity observed experimentally^1,11^ (Fig. 1a). However, such networks exhibit a fundamental trade-off between stability and flexibility^1^. Networks configured to generate stable attractor dynamics are difficult to shut off once activated^5^. Computational models therefore often implement gates to control the timing of attractor dynamics^4,6,7,12^. How such gates might be implemented at a circuit and synaptic level is unclear.

**Figure 1:**
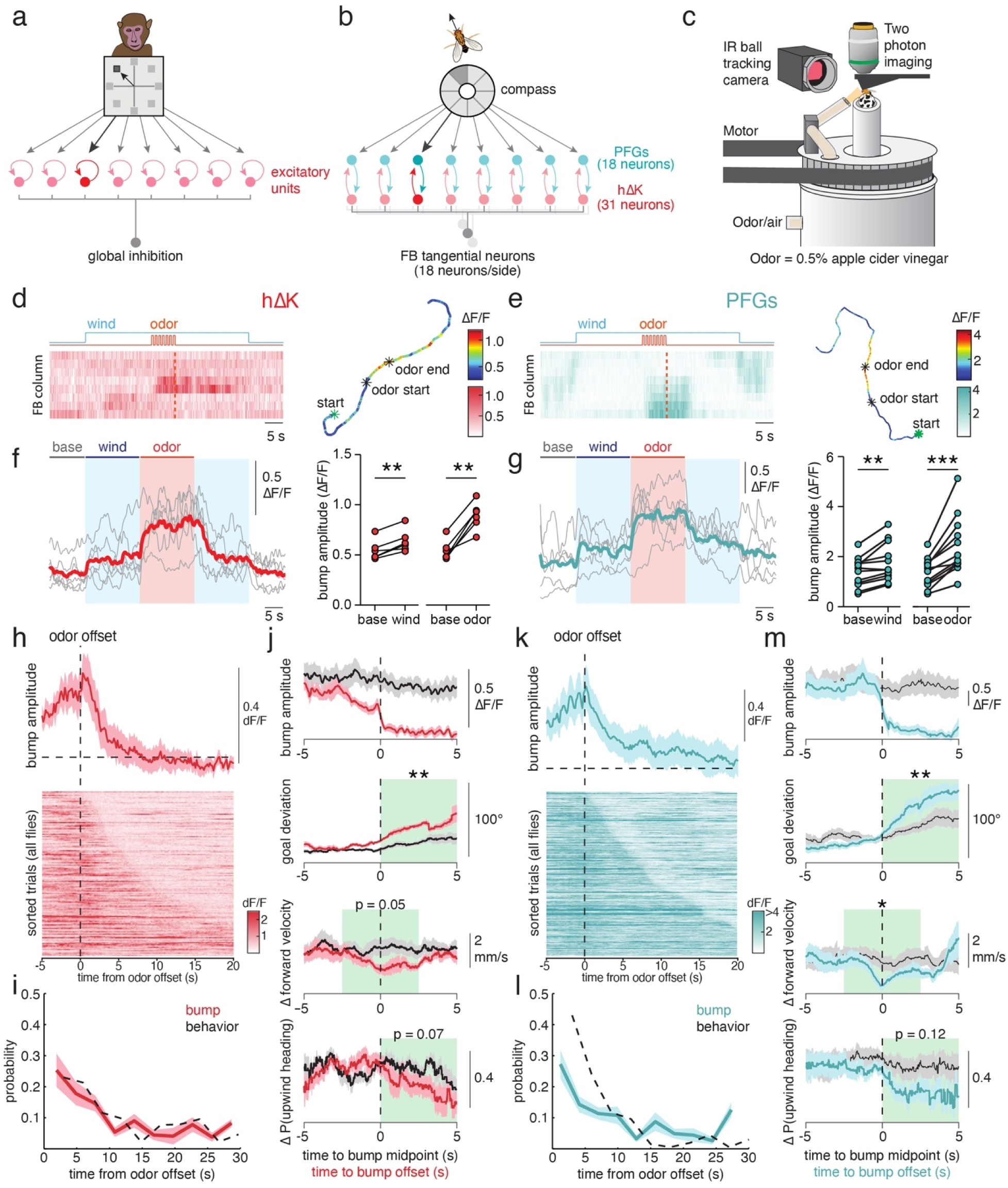
Recurrently connected hΔK and PFG neurons show similar encoding of odor and goal-directed behavior. **a,** Schematic of a recurrent network model developed to describe working memory activity in macaque prefrontal cortex (adapted from Compte et al.^1^). The recurrent network is composed of local recurrent excitation and global inhibition, and is structured to maintain a spatially-tuned direction signal over time. **b,** Schematic of the recurrent network architecture between hΔK and PFG neurons in the FB. hΔK and PFG neurons display local recurrent excitation along the FB, PFG neurons receive spatially structured inputs from the compass, and both hΔK and PFG neurons receive global inputs from ∼18 FB tangential neurons that each provide at least 5% of tangential input to either hΔK or PFG neurons. **c,** Schematic of the *in vivo* imaging paradigm. Fly movement on an air-supported ball is tracked using IR light and an IR-sensitive camera. A tube that can deliver wind and 0.5% ACV rotates around the fly in closed-loop with the fly’s heading. **d,** Example of hΔK bump activity in response to 7 pulses of ACV at 1 Hz. Left: activity shown as raw ΔF/F across FB columns as a function of time. Right: fictive 2D trajectory of the fly color-coded by bump amplitude. Note activity bump that persists after odor offset. **e,** Same as **d** but for PFG neurons. **f,** Mean bump amplitude of hΔK neurons during baseline, wind, and a 15s pulse of ACV. Left: mean activity for each fly (grey) and across flies (red) as a function of time. Right: average activity for each fly as a function of phase. Colored bars on the left plot indicate the windows for averaging for the right plot. **g,** Same as **f** but for PFG neurons. Individual flies in grey and average in blue. **h,** Bump amplitude of hΔK neurons aligned to odor offset for all stimuli. Top: mean bump amplitude aligned to odor offset (error = mean ± SEM); bottom: individual trials sorted by duration of persistent activity. Red dashed line indicates odor offset and black dashed line indicates mean bump amplitude before odor onset. **i,** Distributions of hΔK bump persistence time (red, error = mean ± SEM) and goal directed walking (black) following odor offset. **j,** Mean bump amplitude, goal deviation, change in forward velocity, and change in probability of upwind heading aligned to bump offset (red) or the center of bump periods (grey) (error = mean ± SEM). Green bars indicate the time bin for statistical evaluation. **k-m,** Same as **h-j** but for PFG neurons.

The *Drosophila* brain offers a unique opportunity to study the function of recurrent circuits at a cellular level^13–15^. Connectomics describes the architecture of fly networks with synaptic and cellular precision^8,16,17^, while precise genetic tools enable functional interrogation of the same networks^18^. The fan-shaped body (FB) is a highly recurrent part of the fly navigation center called the Central Complex, thought to compute and encode allocentric goals^8,19–25^. The FB contains ∼30 types of local neurons that interact through complex patterns of feedforward and recurrent connectivity^8^. Several FB local neuron types have been shown to encode goal direction signals for navigation as positionally stable bumps of activity across the columns of the FB^22–24,26–28^. FB output neurons compare these goal signals to an internal compass, computed in the ellipsoid body (EB)^29^ to drive egocentric steering^19,24,25^. How goal signals in local neurons are dynamically regulated to generate moment-to-moment changes in behavior is not clear. The function of recurrent connections between local neuron types also remains unexplored.

Here we investigate how stable activity and rapid switching are combined in a discrete recurrent microcircuit within the FB. When flies navigate towards food, brief encounters with odor drive persistent goal-directed runs that are biased in the upwind direction^26,30^. In a previous study^26^, we showed that a group of FB local neurons called hΔK exhibit a stable bump of activity that turns on with odor and persists while the fly maintains a stable heading after odor loss. Sparse activation of hΔK neurons caused flies to walk in repeatable directions relative to wind^22^, while silencing led to less persistent trajectories after odor loss^26^. Together these data support the idea that bump activity in hΔK neurons represents a persistent goal signal for navigation.

hΔK neurons are embedded in a recurrent network reminiscent of classic models of working memory (Fig. 1a,b), as indicated by three shared motifs. First, hΔK neurons are tightly recurrently connected with a second group of FB local neurons, called PFGs (Fig. 1b; Ext. Data Fig. 1a-c). This recurrent connectivity is highly structured, with strong recurrence within columns of the FB and little connectivity across columns (Ext. Data Fig. 1c,d,f). Second, PFG neurons receive direct inputs from EB compass neurons, which represent heading as a continuously updating bump of activity^13,29^. These inputs are also spatially structured, such that each column of the FB receives heading inputs tuned to different segments of the world (Fig. 1b; Ext. Data Fig. 1d)^8^. Finally, hΔK and PFG neurons both receive global inputs from several types of broadly-projecting tangential neurons (Fig 1b; Ext Data Fig. 1e). The similarity of this network to classic working memory models suggests that this architecture might function to store heading representations over time to orchestrate stable goal-directed walking.

Consistent with the hypothesis that persistent goal representations emerge through hΔK-PFG recurrent interactions, we find that hΔK and PFG show similar persistent bump activity that is activated by odor and correlates with persistent goal-directed runs. We also show that persistent activity in hΔK can be driven by synaptic input but not current injection, and that it requires recurrent signaling from hΔK. We identify inhibitory tangential neurons that could provide the global inhibitory signals necessary for working memory circuit function, and show that excitation and inhibition onto hΔK have distinct dynamics that promote tuneable persistent attractor states across a range of excitation and inhibition weights.

Finally, we examine the mechanisms that allow hΔK activity to be rapidly turned on by odor and off when the fly turns. Surprisingly, we observe that even though hΔK and PFG are tightly anatomically coupled, they only show similar activity during goal directed runs, and their activity diverges during turning and rest. We show in our model that we can reproduce these dynamics by using feedforward inhibition to dynamically decouple hΔK from PFG in a stimulus and behavior-dependent manner. Imaging from inhibitory inputs onto hΔK supports this model, showing that hΔK neurons are inhibited during turning and disinhibited during odor and goal-directed runs. Our data and models reveal a circuit motif— disinhibition of a split recurrent network— that allows stable attractor dynamics to be rapidly switched on and off by either sensory input or ongoing behavior. We propose that this motif may represent a general solution to the problem of combining stability and flexibility in recurrent circuits.

### Recurrently connected hΔK and PFG neurons show similar odor-evoked and goal-associated persistent activity

hΔK and PFG neurons are strongly and reciprocally connected within columns of the FB, with highly correlated forward and reverse connections (Fig. 1b; Ext. Data Fig. 1f,g). Reciprocal connections between hΔK and PFG are the most numerous of any recurrent connection in the Central Complex (Ext. Data Fig. 1h). hΔK neurons express the fast excitatory neurotransmitter acetylcholine, while PFGs express tyramine, MIP, and Dh31, but no classical fast transmitter^18^. To test the hypothesis that these recurrent connections contribute to persistent bump activity previously observed in a broad hΔK line^26^ (VT062617-Gal4), we imaged from each population using cell-type specific split-Gal4 lines^18^. We recorded neural activity during closed-loop odor-and wind-guided navigation^26^ (Fig 1c). In this paradigm, a tethered fly walks on an air-supported ball that controls the orientation of a wind stimulus in closed-loop to simulate wind from a fixed direction. During each 50 second wind trial, we presented odor (0.5% apple cider vinegar) as either a 15 s step, or as 1-10 500ms pulses at 1Hz. In response to odor, flies selected a heading trajectory that was biased upwind (Ext. Data Fig. 1i-k) and persisted for several seconds after odor offset (Ext. Data Fig. 1j,k).

We observed that both hΔK and PFG neurons showed a localized bump of activity that was preferentially activated during odor (Fig. 1d-g, Ext. Data Fig. 2a-c) and persisted after odor offset while the fly maintained its trajectory (Fig. 1h-m; Ext. Data Fig. 2d). In both populations, the probability and amplitude of bump activity grew similarly with pulse number (Ext. Data Fig. 2b,c). In hΔK, the distribution of bump persistence times was very similar to the distribution of goal-directed walking times (Fig. 1i), while in PFG the neural distribution was slightly longer (Fig. 1l), a difference we address in detail below. To ask whether neural persistence predicts behavioral persistence, we detected all bump offsets that occurred after the odor stimulus, and compared trajectories associated with bump offset to those associated with maintained bump activity (Fig 1j,m). In both hΔK and PFG neurons, we found that bump offset was associated with a significant deviation of heading from the ‘goal’ trajectory adopted during odor, compared to periods when bump amplitude was maintained. The presence of bumps was also weakly associated with higher forward velocities, and a greater probability of orienting upwind (Fig. 1j,m). Together, our results argue that hΔK and PFG neurons show similar odor response dynamics and similar persistent activity associated with maintaining an upwind-biased goal heading. Our imaging results thus support the hypothesis that recurrent connectivity between hΔK and PFG contributes to the persistent dynamics observed in both populations during goal-directed running.

### Synaptic contributions to persistent activity in hΔK

We next asked whether persistent activity might arise in part from intrinsic properties of hΔK neurons. We performed whole-cell patch clamp recordings from hΔK neurons (labeled by VT062617-lexA) and injected positive or negative current. We observed rapid response kinetics to current injection (Fig. 2a, Ext. Data Fig. 3a,b) with no persistent excitation. We conclude that intrinsic conductances alone are not sufficient to generate persistent excitation in hΔK.

**Figure 2:**
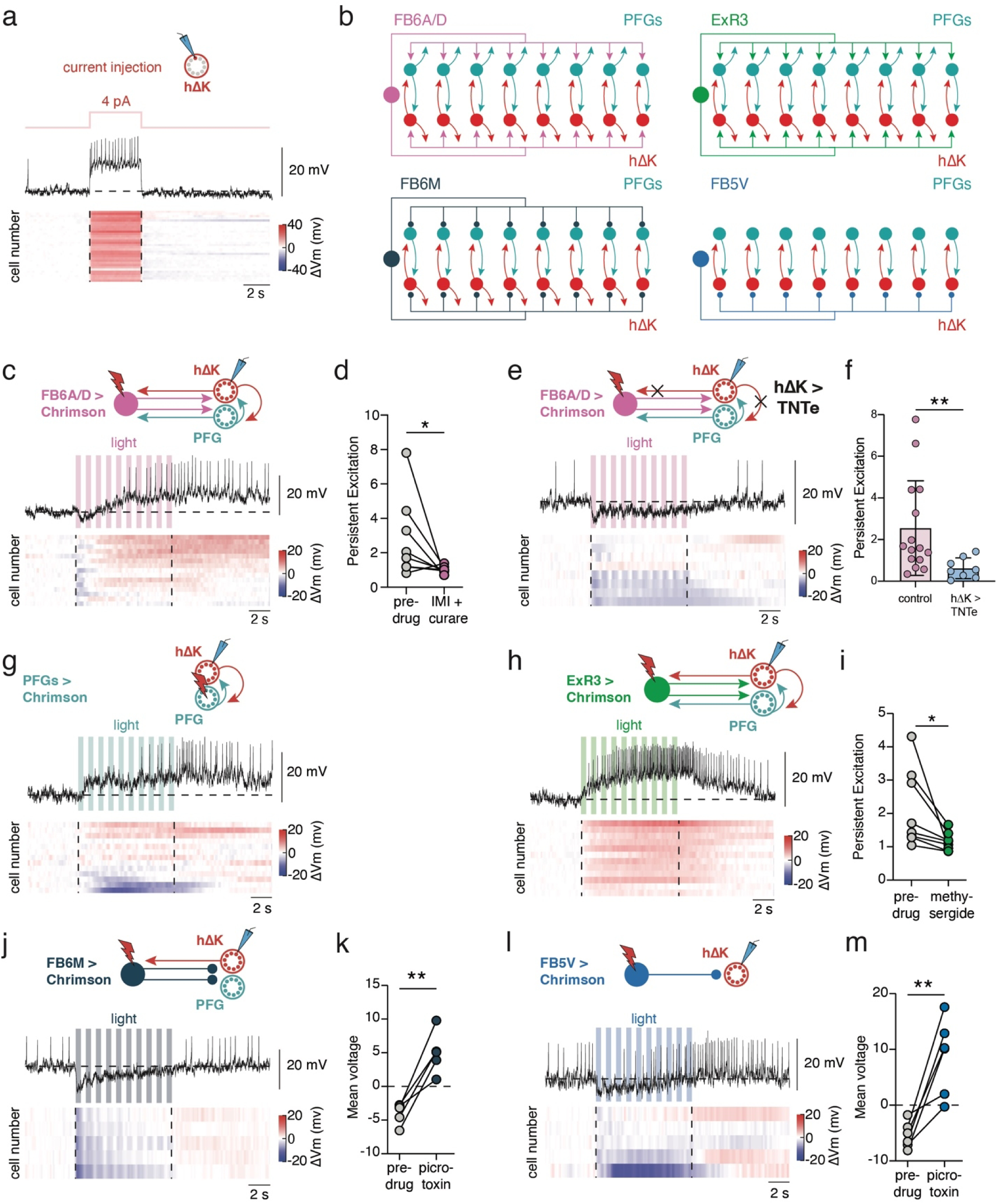
hΔK neurons receive slow excitation and fast inhibition from their tangential partners. **a,** Spiking and membrane potential response of hΔK neurons (VT062617-lexA) during a 4 pA current injection. Top: example trace; bottom: heat map of the mean membrane potential response with each cell as a row. **b,** Schematic of connectivity between tangential inputs (pink: FB6A/D, green: ExR3, dark blue: FB6M, light blue: FB5V) and hΔK and PFG neurons. **c,** Voltage response of hΔK neurons to optogenetic activation of FB6A/D (65C03-Gal4). Same data format as **a**. **d,** Persistent excitation (see Methods) following activation of FB6A/D before and after application of the nicotinic acetylcholine receptor blockers IMI (100 nM) and curare (10 µM). **e,** Spiking and membrane potential response of hΔK neurons following optogenetic activation of FB6A/D when all hΔK neurons express tetanus toxin (TNTe) to block synaptic transmission. Same data format as **a**. **f,** Persistent excitation following activation of FB6A/D in control hΔK neurons (same data as **c**) and neurons expressing TNTe. **g**, Spiking and membrane responses of hΔK neurons following optogenetic activation of all PFG neurons (SS52590 split). Same data format as **a**. **h,** Spiking and membrane potential response of hΔK neurons following optogenetic activation of ExR3 (24C07-gal4). Same data format as **a**. **i,** Persistent excitation following activation of ExR3 before and after application of the general serotonin receptor antagonist methysergide (50 µM). **j,** Spiking and membrane potential response of hΔK neurons following optogenetic activation of FB6M (49B08AD; VT063626DBD). Same data format as **a**. **k,** Mean voltage response of hΔK neurons during optogenetic stimulation of FB6M before and after adding picrotoxin (5 µM). **l,** Spiking and membrane potential responses of hΔK neurons following optogenetic activation of FB5V (VT013602AD; 78H12DBD). Same data format as **a**. **m,** Mean voltage response of hΔK neurons during optogenetic stimulation of FB5V before and after adding picrotoxin (5 µM).

Since current injection did not drive persistent activity in hΔK neurons, we wondered whether synaptic input through tangential neurons could generate persistence. hΔK receives input from several tangential neuron types (Fig. 2b), with different connectivity patterns and predicted neurotransmitters^8,31^. To test the hypothesis that tangential inputs can drive persistent activity, we stimulated a group of tangential neurons that was previously shown to encode odor and drive upwind running^22^ (65C03-Gal4) while recording from hΔK neurons using an orthogonal expression system (VT062617-lexA, Fig. 2c). 65C03-Gal4 labels 17 tangential neurons, at least two of which connected strongly and reciprocally with both hΔK and PFG (FB6A and D, Ext. Data. Fig. 3c). We observed a brief inhibition that switched to slow and persistent excitation later in the stimulus (Fig. 2c). Persistent excitation grew with duty cycle (Ext. Data Fig. 3d,e), and was strongly reduced by the nicotinic antagonists curare (10 µM) and imidacloprid (100 nM, Fig. 2d, Ext. Data Fig. 3f)^32^, arguing that it depends on cholinergic signaling. These neurons have been shown to express both cholinergic markers^18,22^, and markers for glutamate^18^ (an inhibitory transmitter in the fly brain^33^) which might account for the transient inhibition during stimulation. To ask whether recurrent signaling is required for persistent hΔK activity, we stimulated the same tangential neurons, and recorded from hΔK neurons while blocking recurrent signalling from hΔK using tetanus toxin (Fig. 2e). We found that blocking recurrent signaling abolished persistent excitation, and switched the response of hΔK from excitation to weak inhibition (Fig. 2e,f). We conclude that hΔK neurons can generate persistent activity in response to synaptic input, that this persistence requires cholinergic excitation, and that it depends on recurrent signaling by hΔK neurons.

To determine if PFG neurons excite hΔK neurons, we used the same genetic strategy to stimulate PFGs and record from individual hΔK neurons. Upon stimulating PFGs, we observed excitation in some cells and inhibition in others (Fig. 2g), likely because optogenetic stimulation produces an unrealistic depolarization of all members of the class, rather than a physiological bump. hΔK neurons that responded with excitation showed slow and persistent dynamics, supporting the idea that excitatory input from PFGs onto hΔK is also slow. Interestingly, we also observed slow persistent excitation in hΔK when we stimulated a second recurrently connected tangential neuron which expresses serotonin as its major neurotransmitter^34^ (ExR3, Fig. 2b,h; Ext. Data Fig. 3g-h). This persistent excitation was reduced by the broad serotonin receptor antagonist methysergide (50 µM, Fig. 2h,i, Ext. Data Fig. 3i). Together these data suggest that hΔK neurons display persistent excitation in response to multiple recurrently-connected neuron types expressing diverse transmitters and modulators.

In most computational models of recurrent attractor networks, broad inhibition is required together with local recurrent excitation to prevent runaway excitation^1,11,35^. We therefore wondered whether some tangential inputs provide global inhibition onto hΔK. Using connectomic predictions of neurotransmitter phenotype^16,31^, we identified two putative inhibitory tangential neuron types that synapse onto hΔK, called FB5V and FB6M (Fig. 2b), and made or identified lines that label each type (Ext. Data Fig. 4a,b). To test whether these neurons provide functional inhibition onto hΔK, we stimulated each neuron type and recorded from hΔK using the same genetic strategy described above. In all lines we observed strong and rapid inhibition that depressed over time (Fig. 2j,l; Ext. Data Fig. 4c). For both neuron types, stimulus-evoked inhibition was converted to excitation by 5 µM picrotoxin, a concentration known to block GABA-A receptors^33^ (Fig. 2k,m; Ext. Data Fig. 4d). We conclude that some tangential inputs provide broad and rapid inhibition to the hΔK-PFG circuit, consistent with computational models of attractor network design.

### Slow recurrent excitation increases the stability and tuneability of persistent activity in recurrent networks

Our imaging and physiology experiments, together with connectomics, argue that the hΔK-PFG circuit contains all the necessary ingredients to form an attractor network— local recurrent excitation and broad inhibition. However, our experiments also revealed an unexpected finding: synaptic excitation onto hΔK is slow, while inhibition is fast. This slowness may reflect the fact that PFGs as well as many tangential neurons express peptides and modulators in addition to or instead of classical fast neurotransmitters^18^. To understand how these synaptic dynamics contribute to recurrent network function, we developed a connectome-based computational model. In our model, the firing rates of individual hΔK and PFG neurons, and a single global inhibitory tangential neuron, depend on both external input and on recurrent synaptic input between these neurons, as well as a leak term (Fig. 3a, see Methods). The connections between the 30 hΔK and 18 PFG neurons were specified by a Gaussian function fit to the structure of recurrent connectivity between hΔK and PFG observed in the hemibrain connectome^8^ (Fig. 3b, Ext. Data Fig. 5a,b). The tangential neuron receives input from all hΔK and PFG neurons, and provides global inhibition to both populations. The global strength of excitation and inhibition could be scaled through two free parameters (Fig. 3b).

**Figure 3:**
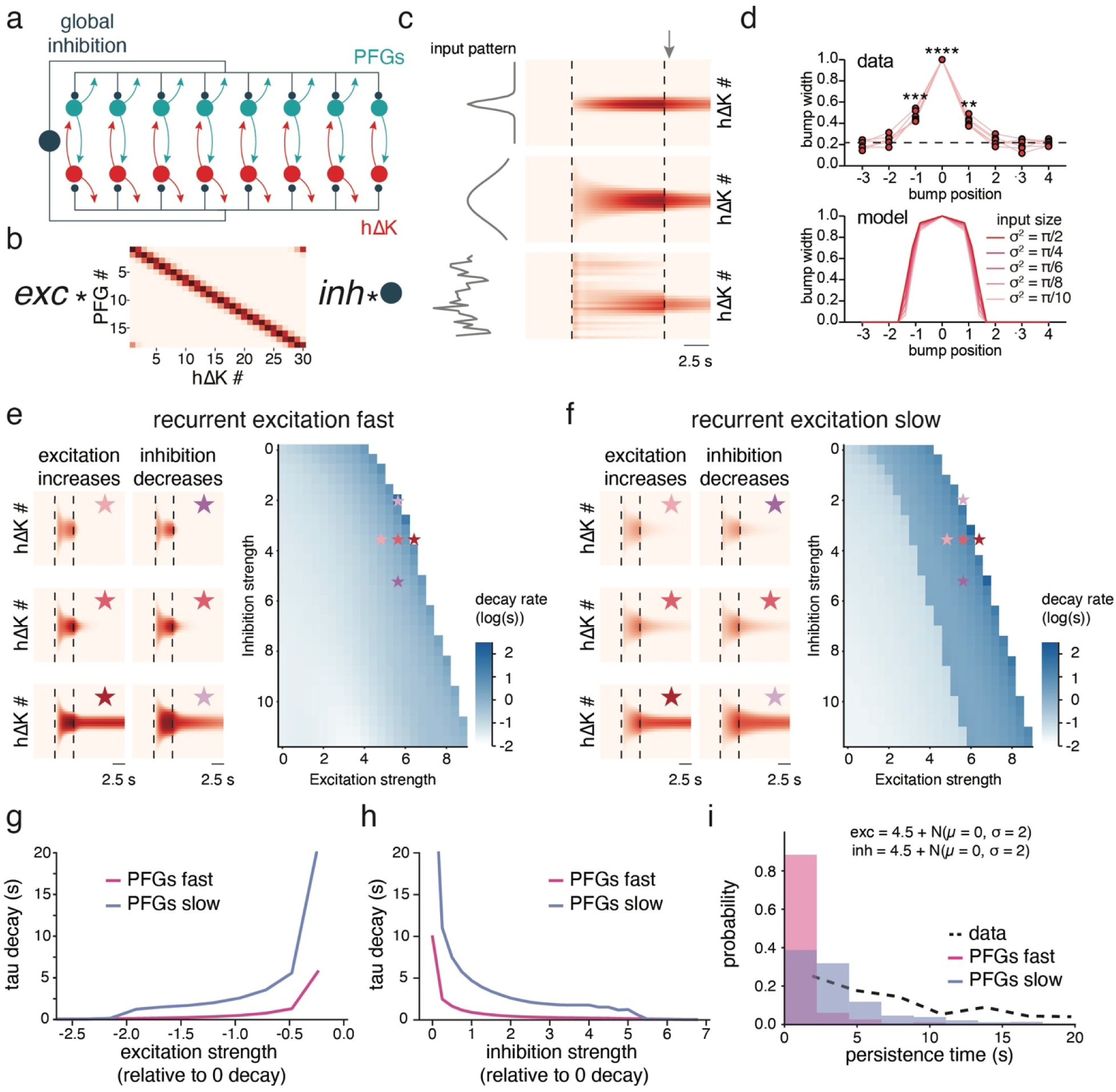
Slow recurrent excitation promotes more stable and tuneable attractor dynamics. **a,** Schematic of connectivity used in simulations. hΔK and PFG neurons are recurrently connected through ring recurrence and both populations are recurrently connected to a global inhibitory neuron. **b,** Schematic showing how the weight matrices are implemented in the model. The weight matrices between hΔK and PFG neurons are scaled by the parameter *exc* (excitation strength), while the connectivity of hΔK and PFG with the global inhibitory neuron are scaled by the parameter *inh* (inhibition strength). **c,** Simulations of hΔK activity in response to narrow Gaussian input (σ^2^ = π/2), wide Gaussian input (σ^2^ = π/10), or random input. Black dashed lines indicate when the inputs are on. Note the similarity in bump width following input offset across all input shapes (arrow). **d,** hΔK bump width from calcium imaging data (top) or estimated from model simulations (bottom, see Methods). **e,** Simulations of hΔK activity using fast recurrent excitation across various excitation and inhibition strengths *exc* and *inh*. Right: decay rate (log_10_ scale) of the network following input offset as a function of *exc* and *inh*. White indicates zero network decay. Left: Example dynamics at various locations in parameter space (stars). **f,** Same as **e** but using slow recurrent excitation in PFG neurons (see Methods). **g,** Mean decay time (tau) of the network as a function of excitation strength using fast (pink) or slow (blue) recurrent excitation. Averages are composed of rows of **e** and **f**, respectively, each aligned to the smallest excitation strength that generated non-decaying persistence. **h,** Mean decay time (tau) of the network as a function of inhibition strength using fast (pink) or slow (blue) recurrent excitation. Averages are composed of the columns of **e** and **f**, respectively, each aligned to the largest inhibition strength that generated non-decaying persistence. **i,** Distributions of persistence times in the fast (pink) and slow (blue) models when the excitation and inhibition strengths are randomly drawn from a Gaussian distribution. Simulations that generated non-decaying persistence were excluded. For comparison, the persistence distribution of hΔK activity following odor offset from Fig. 1i is shown in black.

We first asked whether this network could reproduce the type of persistent bump activity we observed experimentally in hΔK and PFG neurons. We provided spatially structured input to both hΔK and PFG neurons (Fig. 3c) and examined activity after this input was removed. We found that any asymmetric input to the network was transformed into a localized bump of activity with a profile that closely matched those we recorded experimentally (Fig. 3c,d). The width of the attractor bump plateaued exponentially with the width of the Gaussian function describing the local recurrent connectivity (Ext. Data Fig. 5f,g). We also explored whether an attractor bump could be generated with real connectivity values, as continuous attractors that can generate a stable activity bump at any location are difficult to build with noisy biological connectivity^36^. We found that using real connectivity values from the hemibrain connectome^16^, we could generate a bump at most locations in the FB, although bump amplitude and duration varied across locations (Ext. Data Fig. 6a). In contrast, with Gaussian connectivity, a bump of equal magnitude and duration could be generated anywhere around the ring (Ext. Data Fig. 6b). These simulations argue that the recurrent architecture of hΔK, PFG, and inhibitory tangential neurons can support persistent ring attractor dynamics even with real variation in connectivity strength.

Next, we asked how differential synaptic dynamics contribute to network function. To model synaptic dynamics, we filtered the synaptic output of each population by a second-order differential equation to generate an alpha function (see Methods). We fit the two time constants of these alpha functions to either our FB6M activation data for fast synapses (Ext. Data Fig. 5c,e), or to our FB6A/D activation data in the presence of nicotinic receptor blockers for slow synapses (Ext. Data Fig. 5d,e). We used these data because they isolate slow synaptic signaling from FB6A/D by removing acetylcholine signaling from FB6A/D and hΔK neurons. When all synapses were fast, we observed that small changes in global excitation or inhibition strength rapidly transitioned the network from a regime with transient responses (decay < 0.1 s) to a regime with non-decaying responses (Fig 3e). Slow decays similar to those we observed experimentally were only observed for a narrow band of excitation and inhibition weights, implying that these must be fine-tuned to produce realistic decays. In contrast, when recurrent excitation was slow, we observed a large regime of slowly decaying persistence (Fig 3f). Within this regime, small changes in global excitation or inhibition strength gradually modulated the duration of persistence (Fig 3g,h). By incorporating trial-by-trial noise in excitation and inhibition, the slow model, but not the fast model, produced a distribution of decay times approaching the ones we observed experimentally (Fig. 3i). We conclude that the combination of slow excitation and fast inhibition allows the recurrent architecture of hΔK and PFG to produce persistent activity using a broad range of excitation and inhibition strengths, and further allows the duration of persistent activity to be tuned through small changes in these strengths.

Finally we wondered whether the effects of slow excitation on stability and tuneability are specific to the hΔK-PFG circuit architecture, or a more general property of recurrent circuits. To address this question, we generated a simplified recurrent network with one self-recurrent excitatory unit and one bidirectionally connected inhibitory unit (Ext. Data Fig. 7). By assuming that the membrane time constant and synaptic onset kinetics were much faster than synaptic offset kinetics, we created a dynamical system with only two variables: recurrent excitation and inhibition (see Methods). Like our full model, this simple network showed persistent activity that depended on the speed of recurrent excitation: fast excitation led to a narrow band of excitation and inhibition strengths with decaying persistent activity (Ext. Data Fig. 7a), while slow excitation led to a much larger regime of tuneable decay durations (Ext. Data Fig. 7b). By gradually shifting the speed of excitation and inhibition, we found that this regime deteriorates rapidly with excitatory synaptic speed but is relatively insensitive to the kinetics of inhibition (Ext. Data Fig. 7c,d). We conclude that the ability of slow excitation to promote persistence of tuneable duration over a range of excitation and inhibition strengths does not depend on a specific recurrent network configuration.

### hΔK and PFG show can show uncoupled dynamics that depend on behavioral state

Our imaging, electrophysiology, and modeling support the hypothesis that recurrent interactions between hΔK and PFG generate a shared persistent activity bump during odor-evoked goal-directed runs. However, this model does not address how the bump is turned on and off. Indeed, when we simulated the hΔK-PFG circuit described above with a heading input— a continuously varying sinusoidal compass input as suggested by the full connectome of the circuit (Fig. 1b, Ext. Data Fig. 8a)— we observed a bump at most timepoints (Ext. Data Fig. 8a). These observations suggest that some other mechanism must be responsible for turning the bump on with odor and off at the end of a goal-directed run, and led us to examine the dynamics of bump movement in our data more closely.

Close examination of our imaging data revealed that PFG and hΔK neurons showed distinct bump dynamics. In PFGs, a low amplitude bump was present at most times, and translated across columns during turns, becoming brighter during straight goal-directed runs (Fig. 4a). In contrast, hΔK only showed detectable bumps of activity during straight runs, and the bumps that appeared were generally fixed in place (Fig. 4b), consistent with our previous observations^26^. To test whether these differences in bump dynamics held across trials and flies, we measured how far each continuous bump deviated laterally across the FB for both genotypes (Fig. 4c). We found that both the maximum and cumulative lateral deviation were significantly smaller in hΔK compared to PFG (Fig. 4d), supporting the idea that hΔK bumps are generally fixed in position, while PFG bumps translate more readily across columns. To test the hypothesis that bump probability and amplitude depend on behavioral state, we divided trajectories into periods of rest, turns, and straight running (see Methods), and plotted the probability of observing a bump and its average amplitude during each behavior. We found that in both populations, bump probability and bump amplitude were highest during straight running (Fig. 4e,f). During turns and rest, PFGs showed a low amplitude bump, while hΔK rarely showed a bump (Fig. 4e,f). These data argue that hΔK and PFG show similar dynamics mostly during straight goal-directed runs, and that their activity must be functionally uncoupled at other times by some external mechanism.

**Figure 4:**
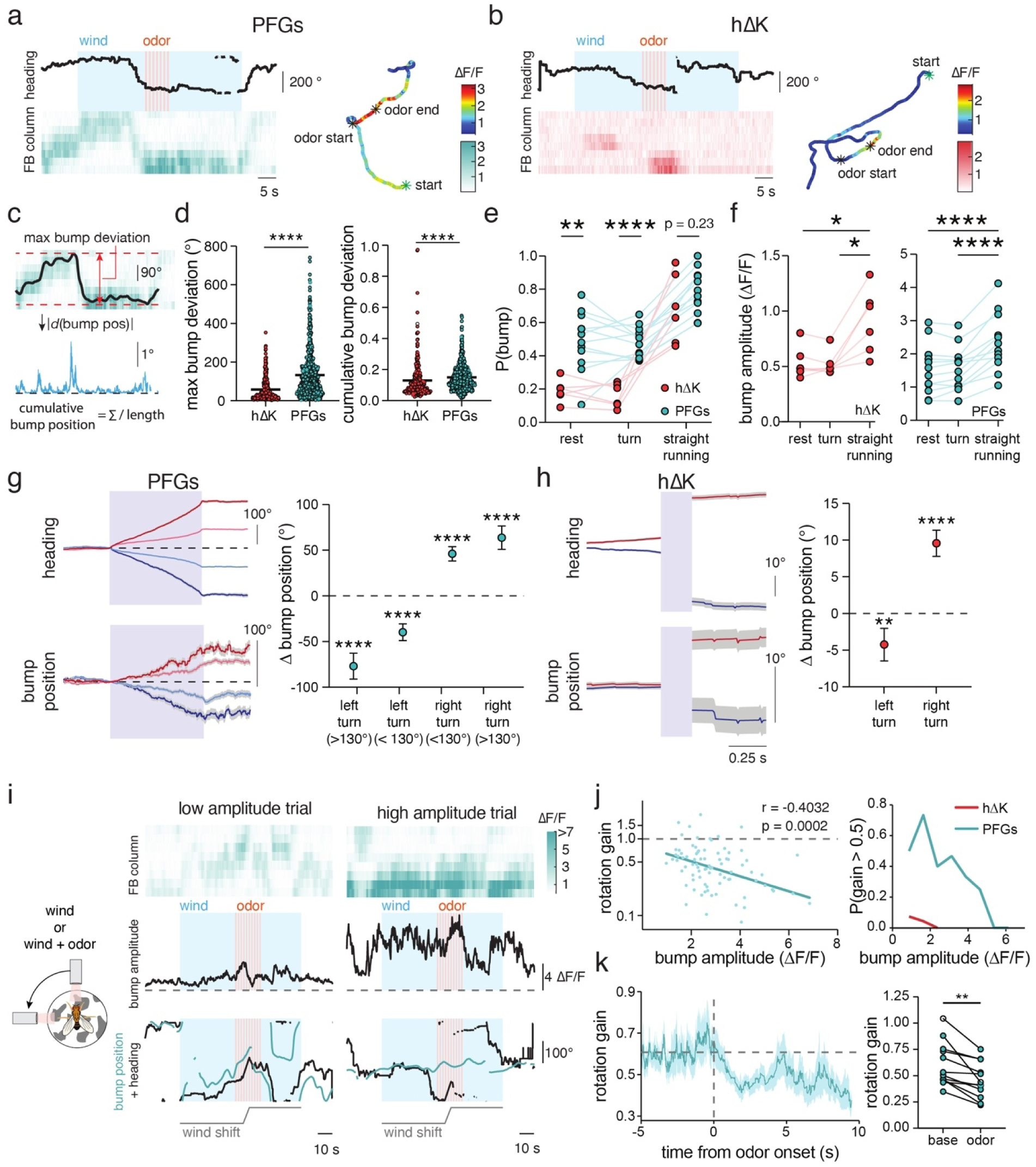
PFG and hΔK can show uncoupled dynamics depending on behavioral state. **a,** Example PFG bump activity in response to 7 pulses of odor at 1 Hz. Left: activity shown as raw ΔF/F across FB columns as a function of time. Right: fictive 2D trajectory of the fly color-coded by bump amplitude. **b,** Same as **a** but for hΔK neurons. Note lateral bump movement in PFG but not in hΔK. **c,** Schematic computation of max and cumulative bump deviation. For max, bump position is unwrapped and maximum difference in bump position is calculated. For cumulative, total change in bump position is summed across time steps and normalized by the bump duration. **d,** Max bump deviation (left) and cumulative bump deviation (right) for all hΔK bumps (red) and PFG bumps (blue). **e,** Bump probability for hΔK (red) and PFG (blue) neurons across rests, turns, and straight runs (see Methods). **f,** Bump amplitude for hΔK (left) and PFG (right) neurons across rests, turns, and straight runs. **g,** Left, mean heading and PFG bump position during large right turns (dark red, Δ heading > 130°), small right turns (light red, 130° > Δ heading > 90°), small left turns (light blue, -90° > Δ heading > -130°), or large left turns (dark blue, -130° < Δ heading). Right, average shifts in PFG bump position across these turns. Positive shifts in bump position correspond with leftward shifts across the FB, and vice versa. **h,** Left, mean heading and hΔK bump position across subsequent bumps separated by right (red, 0° < Δ heading) or left (blue, 0° > Δ heading) turns. Right, average shifts in hΔK bump position across these turns. **i,** Left, schematic for wind shift experiment: wind or wind and odor were shifted by 90° over 2 seconds in a subset of trials. Right, example trials with wind shifts displaying low amplitude PFG bumps (left) or high amplitude PFG bumps (right). Each example shows the raw PFG calcium data (top), bump amplitude (middle), and an overlay of PFG bump position and heading. Note, the wind shift is included in the heading signal. **j**, Left, comparison of the rotation gain and average bump amplitude for PFG bumps present during the wind shifts. Right, fraction of PFG and hΔK data with rotation gains over 0.5 across average bump amplitudes. For hΔK data, analysis was done on non-shift trials, but also see ref^26^. **k**, Rotation gain of PFG neurons over time aligned to odor onset (left) and compared between pre-odor and odor periods (right). Note, this analysis was restricted to non-shift trials.

In many parts of the Central Complex, lateral movement of a bump has been shown to encode a change in heading^13,29^, while in some local neurons, bump movement reflects travelling direction^20,21^. To ask whether lateral shifts in PFG or hΔK bump position encode heading changes, we identified large shifts in heading (between 45-130°), and plotted bump position associated with these turns. We found that PFG bumps reliably shifted across columns during turns, similar to EPG neurons of the heading compass^29^ (Fig. 4g). Such shifts tracked the turning direction, with left turns generating rightward shifts and right turns generating leftward shifts. In contrast, hΔK activity largely disappeared during turns, but could reemerge at a new position following the turn (Fig. 4h). Left and right turns similarly generated opposing direction shifts in hΔK. In single trails, we could observe a positive correlation of PFG bump position with heading (Ext. Data Fig. 8b,d). In contrast, hΔK neurons showed discrete jumps in bump position, with bumps appearing only during periods of relatively fixed heading (Ext. Data Fig. 8c,d). Correlations with travel direction were weaker than those with heading for both populations (Ext. Data Fig. 8e,f), possibly reflecting the lack of optic flow cues in our virtual reality system. We conclude that both PFG and hΔK can track heading changes, but that PFG encodes continuous changes in heading, while hΔK exhibits punctate encoding only during straight runs.

Finally we wondered whether the positionally stable bump activity that we observed during straight runs represents a goal signal (which should not shift with heading^23,24^) or simply a read-out of the fact that the fly is going straight during these periods. In a previous study, we found that hΔK bumps (labeled by the line VT062617-Gal4) did not shift when the wind was rotated during odor, driving compensatory turns, arguing that it represents a goal signal rather than a read-out of straight heading^26^. To ask whether PFG activity can switch between tracking heading and encoding a goal, we performed similar experiments in which we shifted the wind direction by 90° during either wind or odor while imaging from PFG neurons. We found that the behavior of the PFG bump depended on its amplitude (Fig. 4i,j). When PFG bump amplitude was low, bump position shifted with wind rotation and consequent changes in heading (Fig. 4i, left). In contrast, when bump amplitude was high, bump position remained fixed even when the wind was rotated and heading changed (Fig. 4i,right). To quantify this, we computed a rotational gain by comparing how much the bump shifted to how much the heading shifted, where a gain of 1 indicates perfect encoding of heading and a gain of 0 indicates a positionally stable bump. Across bumps, we found that the rotational gain of the PFG bump was negatively correlated with bump amplitude (Fig. 4j, left). Rotational gains above 0.5 were only observed when the PFG bump had a low amplitude, and were almost never observed in hΔK (Fig. 4j, right). We also observed a significant drop in rotational gain following odor onset (Fig. 4k). We conclude that unlike hΔK, whose bump position rarely shifts, PFG can switch from following heading to maintaining a stable goal-like bump position, in a manner that depends on bump amplitude and sensory input.

### Dynamic inhibition of hΔK neurons allows for rapid gating of attractor dynamics

Our imaging results suggest that the hΔK-PFG circuit can operate in two different modes. During goal-directed running, both hΔK and PFG show similar high-amplitude bump activity that is fixed in position, while at other times PFG tracks heading with a low-amplitude bump while hΔK activity is essentially absent. What mechanisms could account for this switch in encoding? To address this question, we revisited the connectome and our computational model. In published connectomes, only PFGs receive direct input from EPG and Δ7 neurons that track fly heading (Fig. 5a, although hΔK does receive input from PEN neurons that encode angular velocity and rotate the heading bump^37, 38^), and this input is significantly weaker than the recurrent connections between hΔK and PFGs (Ext. Data Fig. 9a,b). We wondered whether this difference in input could be combined with gating of recurrent connectivity to produce the dynamics we observed in our imaging. If recurrent connections were shut off, PFGs might follow their compass input, while if recurrence were switched on, their activity would be locked in place at the location of the last compass input. To test this idea, we again implemented a shifting sinusoidal compass input to PFG neurons, and examined activity in the network when recurrent connections between hΔK and PFG were gated OFF or ON (Fig. 5b). Consistent with our prediction, we found that when recurrent excitation was gated OFF, PFG bump position followed the compass signal with low amplitude, while hΔK was silent. In contrast, when recurrent excitation was gated ON, both PFG and hΔK showed a localized high-amplitude bump that remained in place despite shifting input, similar to our experimental observations (Fig. 4a,b,i,j,k).

**Figure 5:**
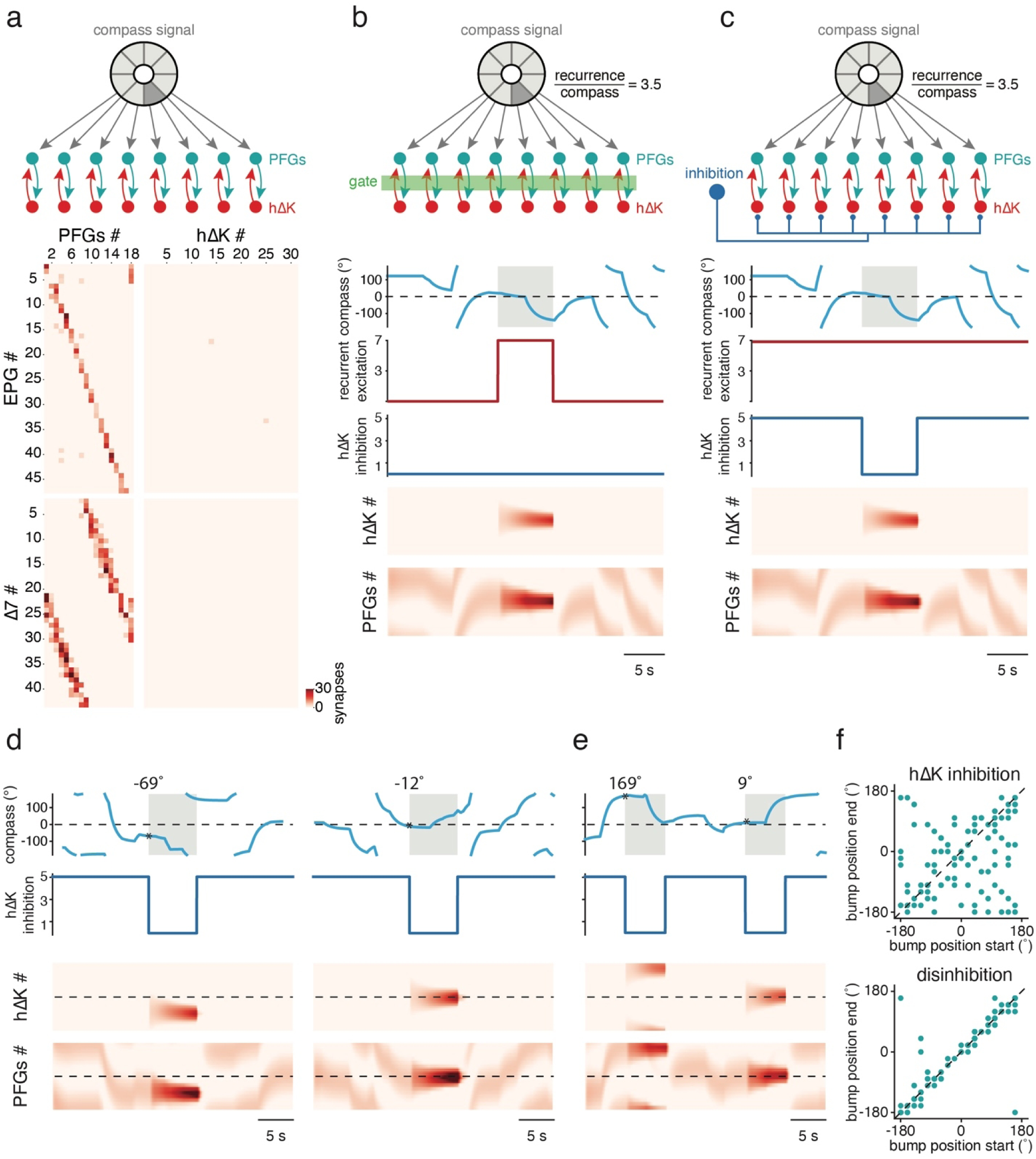
Disinhibition of hΔK allows for rapid gating of recurrent attractor dynamics. **a,** Top, schematic of the connectivity between the compass system of the ellipsoid body and the hΔK-PFG recurrent network. Bottom, synaptic weights from heading-tuned EPG and Δ7 neurons to PFG (left) and hΔK (right) taken from the hemibrain connectome. hΔK and PFG neurons are sorted by their position along the FB, while EPGs are sorted by their position along the EB and Δ7 neurons are sorted by their position along the protocerebral bridge. **b,** Simulations of hΔK and PFG population activity when the EPG compass signal (light blue) is randomly and stochastically updated and recurrent excitation strength (red) is transiently gated from OFF to ON (top). In these simulations, the strength of recurrent synapses relative to compass inputs is 3.5:1. Note higher amplitude stable bump in the presence of recurrent excitation. **c,** Same as **b** but instead of directly modulating recurrent excitation, external global inhibition of hΔK neurons (dark blue) is transiently shut off. Note higher amplitude stable bump during disinhibition. **d,** Further examples of the simulation shown in **c** but where the compass signal at the start of disinhibition varies. Note that the stable hΔK and PFG bump positions depend on these initial compass signals (left: 120°, right: -12°). **e,** Same type of simulation as **c** and **d** but with two bouts of disinhibition separated by a short time interval. Note that each disinhibition bout stabilizes the hΔK and PFG bumps to a different position that depend on the location of the compass input at the time of disinhibition (first: -11°, second: -114°). **f,** Bump position at the beginning and end of a period of hΔK inhibition (top) or disinhibition (bottom) as the compass input randomly fluctuates.

Seeking a more physiological mechanism to rapidly gate connectivity between hΔK and PFG, we wondered whether strong global inhibition of hΔK could serve the same decoupling function. To test this hypothesis, we repeated our simulation but replaced gating of recurrent connections with strong inhibition of hΔK (Fig. 5c). Similar to our results with gated recurrence, we found that when hΔK was inhibited, PFGs followed the compass with a low amplitude bump, while when hΔK was disinhibited, both populations showed a localized high-amplitude bump. Our simulations show that in order for disinhibition to stabilize bump position, recurrent excitation strength must significantly outweigh feedforward heading input (Ext. Data Fig. 9c,d). In multiple published connectomes, the recurrent connections between PFG and hΔK are several-fold stronger than feedforward input from the compass system (Ext. Data Fig. 9b), consistent with the idea that the relative weight of recurrent and feedforward synapses is a reliable feature of PFG wiring.

Finally, we wondered whether this disinhibitory mechanism could be used to rapidly write an ongoing heading signal into memory. We simulated time-varying heading trajectories and removed inhibition when the fly was oriented in different directions (Fig. 5d). This mechanism could be used to rapidly write two different heading memories in quick succession (Fig. 5e). We observed that disinhibition stabilized bump position at the location of the compass input when inhibition was first removed (Fig 5f). We conclude that disinhibition could serve to functionally gate recurrent interactions between anatomically connected partners, rapidly switching the circuit from following an input to storing a pattern in memory.

### Inhibitory inputs to hΔK show signatures of gating

Our model results suggest that disinhibition could support the divergent dynamics we observed in PFG and hΔK neurons, in which PFG can switch from tracking heading at low amplitude, to maintaining a stable bump position at high amplitude, while hΔK bumps are fixed in place and appear only during straight goal-directed runs. If this model is correct, we would expect inhibitory inputs onto hΔK to be tonically active, and to show inhibition during sensory stimuli that promote goal-directed running. We would also expect higher activity during turns, when we observe suppression of hΔK relative to PFG. To test these predictions, we imaged from the two inhibitory tangential neuron types we previously identified through electrophysiology– FB5V and FB6M— as flies performed our olfactory navigation task.

Activity in these inhibitory inputs to hΔK strongly supports our disinhibition model. We first imaged from FB5V neurons (Fig. 6a), a group of 18 neurons (9 per side) that provides feedforward inhibition only onto to hΔK, similar to the inhibitory input we modeled above (Fig. 5c). In these neurons, we observed tonic activity that was strongly supressed during odor, and continued its suppression during maintained trajectories after odor offset (Fig. 6b). Across flies, we observed weak inhibition by wind and strong inhibition by odor (Fig. 6c, Ext. Data Fig. 10a). On a moment-to-moment basis, we found that FB5V activity was negatively correlated with upwind velocity, and positively correlated with angular speed, exactly as predicted by our disinhibition model (Fig. 6d,e). These encoding features might reflect the dendritic inputs to FB5V, which are found in regions encoding both odor (CRE^39^) and pre-motor activity (LAL^40–42^, Fig. 6a). To examine the encoding of turns in more detail, we detected large turns (Δ heading > 130°) and plotted average activity associated with these (Fig. 6f). We observed strongly increasing activity during large heading changes (Fig. 6f,g), consistent with the idea that FB5V inhibits hΔK during large turns. To examine encoding of goal-directed walking, we detected deviations from the goal direction adopted during odor (> 90° from goal direction) and plotted activity associated with these deviations (Fig. 6h). We observed that FB5V activity increased when the fly deviated from its goal (Fig. 6h,i). Our imaging data thus support a role for FB5V suppression in enabling hΔK activity and goal-directed walking, and a role for FB5V elevation in decoupling hΔK from PFG during turns.

**Figure 6:**
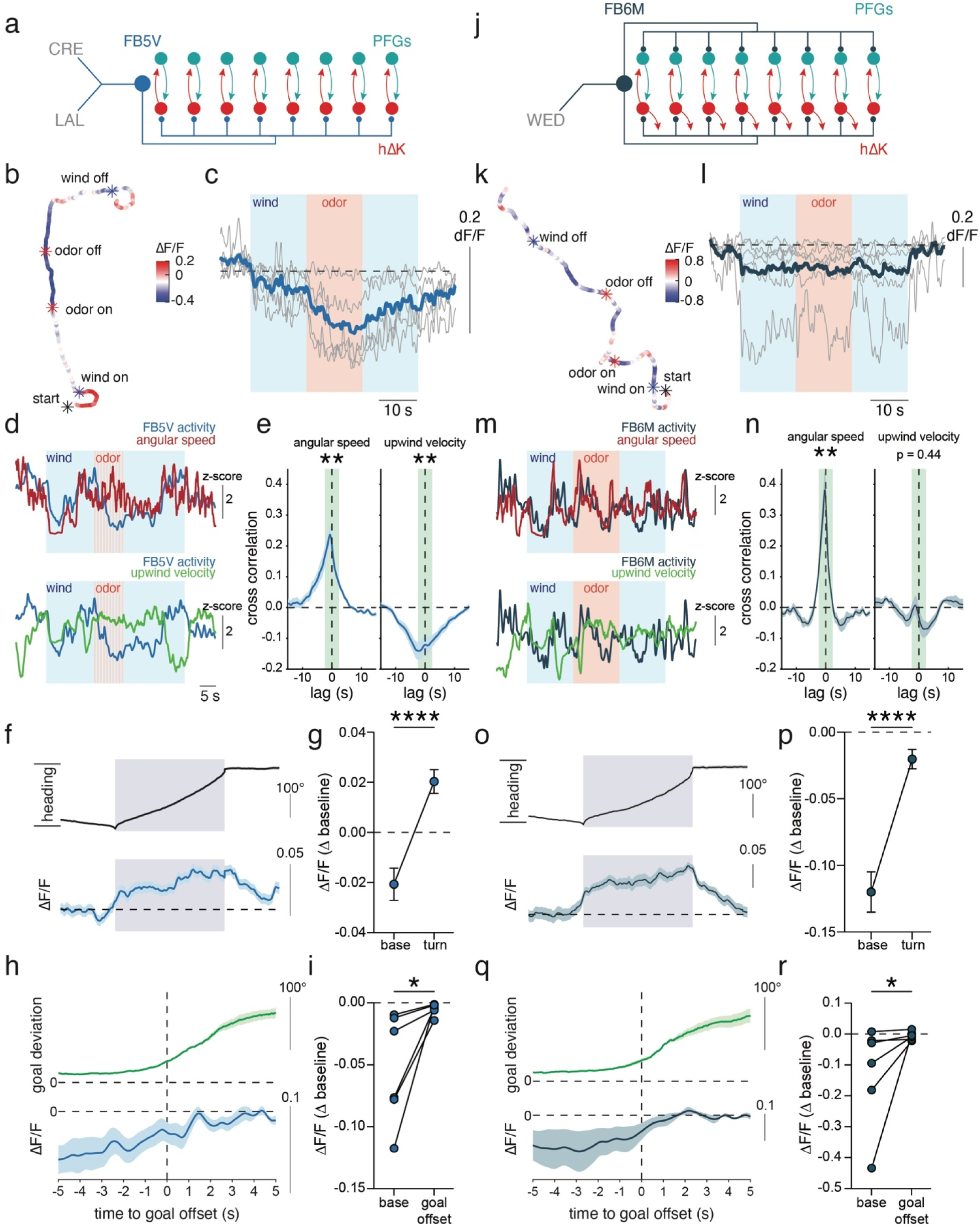
Inhibitory inputs to hΔK show signatures of gating. **a,** Schematic of connectivity between FB5V neurons and the hΔK-PFG recurrent network. Major brain regions innervated by dendrites of FB5V are shown in grey. **b,** Example walking trajectory color coded by activity in FB5V ΔF/F (baseline subtracted). Note inhibition during odor that persists during continued upwind walking. **c,** Mean ΔF/F (baseline subtracted) of FB5V during baseline, wind, and 15s of ACV. Individual flies in grey, average across flies in blue. **d,** Example of FB5V ΔF/F (blue), angular speed (red), and upwind velocity (green) in response to wind and 10 pulses of ACV at 1 Hz. All data z-scored. **e,** Cross correlation between FB5V ΔF/F and angular speed (left) or upwind velocity (right). Statistics correspond to the average cross correlation from -2 to 2 s, shown in green. **f,** Mean heading (absolute value and baseline subtracted) and FB5V ΔF/F during large (>130°) turns. Error = mean ± SEM. **g,** Mean FB5V ΔF/F before and at the end of the turns isolated in **f** (error = mean ± SEM). **h,** Deviation from goal direction and FB5V ΔF/F aligned to the offset of goal-directed walking following odor offset (see Methods, error = mean ± SEM). **i,** Mean FB5V ΔF/F during goal-directed walking (base) and following the offset of goal-directed walking. **j-r,** Same as **a-i**, respectively, but for FB6M.

We also examined the activity of the second tangential neuron type we identified, called FB6M. This group of 4 neurons provides input to both hΔK and PFG and receives feedback from hΔK (Fig. 6j). In this cell type, we also observed tonic activity, with suppression only in response to wind (Fig. 6k,l, Ext. Data. Fig. 10b). FB6M activity during wind was positively correlated with angular speed, but showed no correlation with upwind velocity (Fig. 6m,n). Like FB5V, FB6M showed increased activity during turns (Fig. 6o,p) and when the fly deviated from its goal direction (Fig 6q,r). The wind encoding of FB6M might reflect its inputs in mechanosensory regions, which are known to encode the presence of wind (WED^43,44^, Fig. 6j). In our model, we find that inhibition of both hΔK and PFG does not decouple their activity but instead shuts off both populations (Ext. Data Fig. 9e). We conclude that FB6M also plays a disinhibitory role in enabling circuit activity during wind and straight walking, but may permissively regulate the activity of PFG and hΔK together, rather than serving to decouple these populations.

## Discussion

Attractor networks composed of recurrently connected neurons contribute to many neural functions, including gaze stabilization^45–46^, spatial working memory^9^, motor control and decision making^10^, navigation^13,47^, and emotional states^48,49^. By forcing diverse patterns of input onto a limited set of stable states, attractor networks offer many advantages for computation^2^, such as de-noising^50^, evidence integration^51,52^, and short-term information storage^1^. A major issue in building attractor networks is that network architecture alone cannot always produce the desired dynamics. In particular, recurrent networks with realistic fast units often fail to reproduce the kind of slow persistent activity dynamics observed in experiments^35,48^. A second issue is that attractor networks with fixed weights do not generate intrinsic dynamics that are restricted to particular behaviors, sensory inputs, or internal states^10,49^. How biological recurrent networks are structured to generate stable dynamics that can be rapidly switched on and off by sensory input or behavior is unclear.

One solution to generate robust slow dynamics is to make recurrent excitation slow^35,48^. The most common mechanism proposed for generating slow excitation is through NMDA receptors at excitatory glutamatergic synapses^35^, although a variety of other slow biophysical mechanisms, such as synaptic facilitation^51^ and peptide signaling^48^ have also been proposed to play a role in stabilizing recurrent activity. Conversely, to control the onset and offset of attractor dynamics, many network models implement gates to globally scale the strength of recurrent connectivity^4,6,7^. A common motif for regulating the gain of neural activity in biological systems is disinhibition. In cortical circuits, a canonical disinhibitory circuit from VIP interneurons to PV interneurons to pyramidal neurons has been shown to regulate neuronal gain^54^, contrast sensitivity^55^, and learning^56^. In the basal ganglia, brief periods of disinhibition allow for the performance of skilled motor actions^57,58^. Disinhibition also gates visual responses in the retina^59^, and both auditory responses and motor production in songbirds^60,61^. Despite the prevalence of disinhibition, the interaction of disinhibitory gating mechanisms with recurrent circuits has not been investigated in detail.

Here we took advantage of the complete connectome information available in the fly^8,16^, combined with precise genetic access to neuronal populations^18^, to investigate the relationship between anatomical connectivity and functional dynamics in a discrete recurrent circuit in the fly navigation center. Consistent with this circuit using reverberant excitation, we observed similar dynamics in both hΔK and PFG neurons in response to odor and during goal-directed behavior. We found that hΔK neurons exhibit persistent activity in response to synaptic input but not current injection, and we found that blockade of recurrent signaling in hΔK neurons abolished persistent activity. Computational modeling based on the connectome supports the idea that this circuit behaves as a ring attractor, capable of stabilizing activity at any point on the ring, even with the “noisy” connectivity observed in real connectomes, although some of the persistence we observed *in vivo* may also arise upstream, as inhibition in FB5V inputs to hΔK also outlasted the odor stimulus. A recent study identified MBONs upstream of FB5V that modulate movement timing following odors and might contribute to dynamics we observed in FB5V^62^. The architecture of this network bears a striking resemblance to classical models proposed to underlie working memory in the prefrontal cortex^1,63^. However, the accessibility of the fly circuit also reveals several design features that enable this circuit to combine stable representations with rapid gating.

First, slow excitation in the hΔK-PFG circuit appears to be achieved through multiple biophysical mechanisms. PFG neurons do not express any fast transmitter, and instead express only neuromodulators and peptides^18^. Consistent with this expression profile, when we observed excitation in hΔK on stimulating PFGs, this excitation was slow and persistent (Fig. 2g). Our model indicates that slow recurrent excitation from local PFG neurons can generate attractor dynamics over a range of excitation and inhibition strengths and can make the duration of persistent activity tuneable. Our work expands on the classical idea that NMDA receptors are critical for building a stable working memory circuit^1,35^, suggesting that any slow signaling mechanism can serve to stabilize persistent activity. Our results are also consistent with recent findings in the hypothalamus arguing that slow signaling through neuropeptide receptors is required for a persistent aggressive state in mice^46,64,65^.

Second, the recurrent network is split into two parts composed of distinct neuron types: hΔK and PFG. In most attractor models, recurrent excitation occurs within a uniform population of excitatory neurons, parameterized by stimulus tuning, timing, or other features^1,11^. Here instead we find that the network is composed of two morphologically^8^ and transcriptomically^18,66^ distinct neuron types that receive distinct inputs. PFG neurons receive a heading direction signal from the compass system and form the content of memory, while hΔK neurons receive an inhibitory gating signal from FB5V and control memory formation. By separating the input of the network from the gating signal, we find in our models that we can rapidly gate recurrent activity on and off. hΔK and PFG also receive common input, for example from hΔB neurons that encode travel direction^20,21^. Although we did not observe encoding of travel direction in our paradigm, common input might allow this circuit to perform additional computations important for navigation. The separation of the circuit into two halves may also facilitate proper wiring of the network, which requires a high ratio of recurrent to feedforward inputs in order to generate inhibition-dependent switching behavior. Future studies may investigate the molecular signals that allow hΔK and PFG to wire together in a ring structure, that establish the proper ratio between recurrent and feedforward synapses, and that allow hΔK and PFG to wire to both distinct and common inputs.

Finally we show evidence that inhibition of hΔK allows the circuit to rapidly switch between two modes: information following and information storage. Our model circuit functions similar to a latch in digital logic^67^: during inhibition PFGs follow their compass inputs, while during disinhibition activity is locked into place. In our paradigm, we observed that inhibition onto hΔK was supressed by odor and wind— two cues that promote straight upwind trajectories— and that activity of the circuit correlates with upwind-biased orientations. However, in principle this circuit could be used to store any heading direction, as long as the gate is turned on (i.e. inhibition turned off) under the correct conditions. In addition to suppression by sensory input, we also observed that inhibitory inputs onto hΔK are active during turning, suggesting that the circuit can be gated by motor features as well as sensory ones. In the future, it will be important to rigorously test our disinhibition model by imaging from hΔK or PFG neurons while manipulating FB5V activity.

The mechanism described here is reminiscent of landmark learning in the compass system, where dopaminergic plasticity is gated by turning^68^. However, disinhibition allows for near instantaneous— though short-lived— memory formation. Several previous models have used asymmetric synapses between tangential neurons and FB local neurons to store directional memories^27,69,70^, although such a mechanism has not yet been demonstrated experimentally. A variety of plasticity mechanisms, operating at different timescales and gated by diverse sensory and motor features, are likely to underlie the complex dynamics of biological goal signals in the fly navigation center.

Gated recurrent networks have been theorized to initiate intrinsic dynamics, such as persistent activity, and to coordinate more complex forms of dynamics, such as the manipulation of information in working memory and motor preparation and control^3,6^. Similar types of gates are used extensively in machine learning^71^, including as a solution to rapidly reset memory traces^72^. Here we describe a biological form of gating that allows a recurrent anatomical circuit to dynamically alter its function on fast behavioral timescales. If integrated into recurrent networks with different connectivity structures^2,73^, disinhibition could be used to rapidly alter the function of other types of attractor networks, allowing for flexible use of line attractors, rotations, sequences, or bistable attractors in a stimulus- and behavior-dependent manner.

## Methods

### Fly stocks

All experimental flies were raised on standard cornmeal-agar media at 25 °C on a 12-hour light cycle. For optogenetics experiments, the food was supplemented with 50 µL of 35 mM all-trans retinal (Sigma, R2500, dissolved in ethanol) mixed with ∼1 teaspoon of hydrated potato flakes. Experimental flies were aged to 1-3 days for electrophysiology experiments or 9-14 days for imaging experiments. A list of fly genotypes is shown in **Table 1**.

**Table 1:** Fly genotypes.

| Genotype | Figures | Notes |
| --- | --- | --- |
| VT062617AD; VT037223DBD > UAS-GCaMP7f, UAS-tdTomato | Fig. 1d,f,h,i,j<br>Fig. 4b,d,e,f,h,j<br>Ext. Data Fig. 1i,j,k<br>Ext. Data Fig. 2b,d<br>Ext. Data Fig. 8c,d,e,f | Split Gal4 SS63158 targeting hΔK |
| SS52590 (VT017491AD; VT000355DBD) > UAS-GCaMP7f, UAS-tdTomato | Fig. 1e,g,k,l,m<br>Fig. 4a,c,d,e,f,g,i,j,k<br>Ext. Data Fig. 1j,k<br>Ext. Data Fig. 2c,d<br>Ext. Data Fig. 8b,d,e,f | PFG split (Bl: #86628) |
| VT062617-gal4 > UAS-GCaMP7f, UAS-tdTomato | Ext. Data Fig. 1j,k | Broad line targeting hΔK<sup>26</sup> |
| VT062617-lexA, 13xlexAop-GFP/+; UAS-Chrimson/65C03-gal4 | Fig. 2a,c,d,f<br>Ext. Data Fig. 3a,b,d,e,f<br>Ext. Data Fig. 5d | Recording from hΔK; activate 65C03 (Bl: 41290) is a FB tangential line that drives upwind walking<sup>22</sup> |
| VT062617-lexA, 13xlexAop-GFP/lexAop-TNTE; UAS-Chrimson/65C03-gal4 | Fig. 2e,f | Same as above with TNTE to block synaptic transmission from hΔK |
| VT062617-lexA, 13xlexAop-GFP/ VT017491AD; UAS-Chrimson/ VT000355DBD | Fig. 2a,g<br>Ext. Data Fig. 3a,b | Recording from hΔK; activate PFG split (Bl: #86628) |
| VT062617-lexA, 13xlexAop-GFP/+; UAS-Chrimson/24C07-gal4 | Fig. 2a,h,i<br>Ext. Data Fig. 3a,b,g,h,i | Recording from hΔK; activate ExR3 using 24C07 |
| VT062617-lexA, 13xlexAop-GFP/49B08AD; UAS-Chrimson/VT063626DBD | Fig. 2a,j,k<br>Ext. Data Fig. 3a,b<br>Ext. Data Fig. 5c | Recording from hΔK; activate FB6M using 49B08AD; VT063626DBD |
| VT062617-lexA, 13xlexAop-GFP/VT013602AD; UAS-Chrimson/78H12DBD | Fig. 2l,m | Recording from hΔK; activate FB5V using VT013602AD; 78H12DBD (SS88825) |
| VT013602AD; 78H12DBD > GFP | Ext. Data Fig. 4a | Visualization of split used to target FB5V for activation |
| 36B06-Gal4 > GFP | Ext. Data Fig. 4a | Visualization of broad line used to target FB5V for activation |
| 49B08AD; VT063626DBD > GFP | Ext. Data Fig. 4b | Visualization of split used to target FB6M for activation |
| VT062617-lexA, 13xlexAop-GFP/+; UAS-Chrimson/36B06-gal4 | Ext. Data Fig. 3a,b<br>Ext. Data Fig. 4c,d | Recording from hΔK; activate 36B06 (BI: 49929) targeting FB5V |
| VT062617-lexA, 13xlexAop-GFP/+; UAS-Chrimson/21D07-gal4 | Fig. 2a<br>Ext. Data Fig. 3a,b | Recording from hΔK; activate 21D07 (BI: 48943) targeting FB5AB<sup>22</sup>. Only used for current injections in this paper. |
| VT013602AD; 78H12DBD > UAS-GCaMP7f, UAS-tdTomato | Fig. 6b,c,d,e,f,g,h,i<br>Ext. Data Fig. 10a | Split Gal-4 line SS88825 targeting FB5V |
| 49B08AD; VT063626DBD > UAS-GCaMP7f, UAS-tdTomato | Fig. 6k,l,m,n,o,p,q,r<br>Ext. Data Fig. 10b | Split Gal4 line targeting FB6M:49B08AD; VT063626DBD |

\*BI refers to Bloomington Drosophila Stock Center

### Two-photon calcium imaging

Calcium imaging was performed as described in our previous study^26^. Briefly, we anesthetized flies over ice and a chilled (4-10 °C) aluminum sarcophagus and then glued them onto custom 3D-printed holders. We minimized movement of the brain by stabilizing the proboscis and head, and glueing the eyes, head, and anterior tip of the thorax to the holder. To access the FB for imaging, we removed a small piece of the surface cuticle, trachea, and air sacs over the back of the brain using sharpened forceps. We positioned flies over an air-supported ball (9 mm diameter General plastics FR-7110, painted with black spots, supported by 0.4 L/s air) using two cameras (30fps, FLIR Blackfly, Computar 8.5mm 1:1.3 adjustable lens) at the front and side of the fly. A third camera (100 fps, Grasshopper, 94mm/0.5x InfiniStix Proximity Series lens, Edmund Optics Focal Length Extenders, working distance=15cm) was used to track the ball using Fictrac software^77^. We illuminated the fly and ball using two IR LEDs (M850F2) and T-cube LED Drivers (LEDD1B) coupled to fiber optic cables (Thorlabs M118L03 flat cleave patch cables). We continuously perfused (1mL/min) the brain with extracellular saline (103 mM NaCl, 3 mM KCl, 5 mM TES, 8 mM trehalose dihydrate, 10 mM glucose, 26 mM NaHCO_3_, 1 mM NaH_2_PO_4_H_2_0,1.5 mM CaCl_2_2H_2_O, and 4 mM gCl_2_6H_2_O, pH 7.1–7.4, osmolarity 270–274 mOsm) bubbled with carbogen (5% CO_2_, 95% O_2_) and warmed to 33°C.

We performed all imaging under a 20x water objective (Olympus Plan Fluorite XLUMP 20x/1.0) using an infrared laser (SpectraPhysics MaiTai HP or Coherent Axon 920) at 920 nM and 25-35 mW (measured at sample) using a galvo-galvo scanner. Simultaneous fluorescence emission from GCaMP7f and tdTomato were separated by bandpass filters (Semrock, FF01-525/50-32 for green and FF01-607/70-32 for red) and detected using two GaAsP photomultiplier tubes. We captured volumes consisting of three 192 x 96 µm optical sections, separated by 15 µm, taken at 8 vol/s with a 1.2 µs dwell time.

In all imaging experiments, flies walked on a floating ball and received wind (25 cm/s) and odor (0.5% apple cider vinegar, 0.3 L/min) stimuli controlled by custom Python code. The wind direction was set by a rotary union (Dynamic Sealing Technologies, # LT-2141-OF-ES12-F1-F2-C2) controlled by a stepper motor (Oriental PKP566FMN24A, with controller CVD524-K) and microcontroller (TeensyDuino +USB), and was controlled in closed loop by the flies’ heading measured by ball movement as we described previously^26^. The starting wind direction was randomly selected from 0°, 45°, -45°, 135°, or -135°. Wind was on from 5-60 seconds of each 65 second trial. The odor signal was randomly selected between five possible patterns on interleaved trials: odor step (15 s) or odor pulses of 1, 4, 7, or 10 pulses (1 Hz, 0.5 s pulse width). When the odor was turned off, a compensatory air valve was opened to maintain total air speed. Air to all lines (wind, ball support, odor, compensated air) were passed through a pressure regulator (Cole-Parmer MIR2NA) and charcoal filters (Drierite 26800 Drying column, with desiccant replaced by activated charcoal). Wind and compensated air were also humidified. Airflow for the wind line was controlled by a mass flow controller (Aalborg Model GFC17), while airflow for odor, compensated air, and ball support were controlled by flowmeters (Cole-Parmer PMK1-010608). We controlled the timing of wind stimuli using a solenoid valve (Cole-Parmer Masterflex P/N 01540-11), and the timing of odor and compensate using high-speed three-way solenoid valves (Lee company, LHDA1233115HA).

During wind-shift experiments, the wind was rotated by 90° across 2.2 s either in the presence of odor (10 pulses at 1 Hz) or without odor. The onset of the shift was set to occur 4 s after odor onset. We also included control trials where wind was not shifted in the presence or absence of odor. The data from non-shift control trials were included in all analyses that did not depend on the precise odor stimulus.

### Electrophysiology

For electrophysiology, we anesthetized flies over ice, glued them to custom 3D printed holders, and then removed their two front legs. To access dorsal cell bodies, we removed a large piece of the cuticle, trachea, and air sacs from the back of the brain, then manually detached the sheath using fine forceps. We minimized brain movements by removing the muscles on the brain and stabilizing the proboscis with glue. Starting at the dissection, and maintained throughout the entire experiment, we continuously perfused extracellular saline (same recipe as above), bubbled with carbogen, over the brain. Using a 40× objective (Olympus, LUMPLFLN40XW), an LED source (Cairn Research MONOLED), and a filter (U-N19002 AT-GFP/F LP C164404) we visualized GFP-positive cell bodies for recordings. Prior to recording, we cleaned the area around the target cell using fine-tip glass pipettes filled with extracellular saline.

For patch clamp recordings, we pulled glass pipettes with a Sutter P-1000 puller, pressure polished the pipettes (final resistance of 3-5 MΩ), and then filled them with intracellular solution (140 mM KOH, 140 mM aspartic acid, 10 mM HEPES, 1 mM EGTA, 1 mM KCl, 4 mM MgATP, 0.5 mM Na_3_GTP, and 13 mM biocytin hydrazide). We amplified the voltage signals using an Axonpatch 200B amplifier with a Brownlee Precision 410 preamplifier and then digitized the signals at 10 kHz. We delivered 73 µW/mm^2^ red light (625 nm, measured at light source) for activating CsChrimson using a Thorlabs LED (M625F2) and T-Cube LED Driver (LEDD1B) coupled to optical fibers cables (Thorlabs M118L03 flat cleave patch cables) positioned below the head.

To better observe a mixture of excitation and inhibition, we depolarized cells to -38.1 ± 7.1 mV (mean + SD, N = 37 cells) using a small amount of positive current. Cells that did not spike were discarded. In all experiments, we randomly looped through 11 stimuli: seven optogenetic light stimuli (4 s of 12 µW/mm^2^ low power; 4 s of 23 µM/mm^2^ medium power; 4 s of 73 µM/mm^2^ high power; 1 Hz pulses with 0.2 duty cycle; 1 Hz pulses with 0.5 duty cycle; 1 Hz pulses at 0.8 duty cycle; and a light plume stimulus), three current stimuli (4 s of -4 pA; 4 s of 0 pA; and 4 s of 4 pA), and one wind/odor stimulus (wind on for 20 s, 5% apple cider vinegar on for 4 s). We constructed the light plume stimulus by filtering the odor encounters of a fly navigating in a virtual olfactory plume^26^ with a positive derivative filter. The stimuli were looped five times in all cells, and then five times per drug condition. While we only report the results of light pulse stimuli in this paper, the voltage changes we observed – such as fast inhibition or slow persistent excitation - were seen across all optogenetic temporal patterns used.

In several experiments, we perfused drugs onto the brain to block neurotransmitter receptors. In these experiments, drugs were mixed with extracellular saline and perfused onto the brain for at least five minutes before resuming stimulus presentation. Since picrotoxin generated large excitatory responses to synaptic input, we often hyperpolarized the cells to -50.3 ± 10.2 (mean + SD, N = 10 cells) to minimize depolarization block. If cells did enter depolarization block, we paused the experiment until the cell recovered. The list of drugs, their sources, and the final concentrations used are listed in **Table 2**.

**Table 2:** Fly Reagents.

| Reagent | Source | Identifier |
| --- | --- | --- |
| 20XUAS-IVS-jGCaMP7f (II) | BDSC | RRID:BDSC_80906 |
| UAS-tdTom (II) | BDSC | RRID:BDSC_36327 |
| 13XLexAop-GFP (II) | BDSC | RRID:BDSC_52265 |
| 20XUAS-IVS-CsChrimson.mVenus (III) | BDSC | RRID:BDSC_55136 |
| LexAop-TNT (269b) (II) | Janelia; Sen et al., 2019<sup>74</sup> | N/A |
| 10XUAS-IVS-mCD8::GFP (III) | BDSC | RRID:BDSC_32187 |
| yw;<br>B3RT-vGlut-LexA/CyO,EYFP;<br>UAS-B3,<br>13XLexAop-6XmCherry,<br>20XUAS-6XGFP/TM6c | Steven Stowers, Sherer et al., 2020<sup>75</sup> | N/A |
| SS63158 (VT062617AD; VT037223DBD) | Janelia, gift of Tanya Wolff<sup>18</sup> | Robot ID: 3044006 |
| SS52590 (VT017491AD; VT000355DBD) | BDSC | RRID:BDSC_86628 |
| VT062617-Gal4 (III) | VDRC | VDRC: 206875* |
| VT062617-lexA (II) | Hamid et al., 2024<sup>76</sup> | N/A |
| 65C03-Gal4 (III) | BDSC | RRID:BDSC_41290 |
| 24C07-Gal4 (III) | BDSC | RRID:BDSC_49074 |
| 49B08AD (II) | BDSC | RRID:BDSC_86664 |
| VT063626DBD (III) | BDSC | RRID:BDSC_73940 |
| SS88825 (VT013602AD; 78H12DBD) | Janelia | Robot ID: 3046075 |
| 36B06-Gal4 (III) | BDSC | RRID:BDSC_49929 |
| 21D07-Gal4 (III) | BDSC | RRID:BDSC_48943 |

\*no longer available

### Immunohistochemistry

To perform immunohistochemistry, dissected brains were fixed in 4% paraformaldehyde (dissolved in PBS) for 15 minutes, washed 3x in PBS, incubated in 5% normal goat serum (dissolved in PBST) for 60 minutes, incubated overnight in primary antibody solution (dissolved in 5% normal goat serum), washed 3x in PBST, incubated overnight in secondary antibody solution (dissolved in 5% normal goat serum), washed 3x in PBST, and washed 3x in PBS. The primary antibody solution contained chicken anti-GFP (Fisher Scientific RRID:AB_1074893) 1:50, mouse anti-nc82 (DSHB RRID:AB_2314866) 1:50, and rabbit anti-dsRed (Clontech 632496) 1:500. The secondary antibody solution contained Alexa488-conjugated goat anti-chicken (Fisher Scientific RRID:AB_2534096) 1:250, Alexa633-conjugated goat anti-mouse (Fisher Scientific RRID:AB_2535719) 1:250, and Alexa568-conjugated goat anti-rabbit (Fisher Scientific RRID:AB_2576217) 1:250. For imaging, the brains were mounted on microscope slides, immersed in vectashield (Vector Labs H- 1000), sealed with coverslips, and then imaged using a 20× objective (Zeiss W Plan-Apochromat 20×/1.0 DIC CG 0.17 M27 75 mm) on a Zeiss LSM 800 confocal microscope at 1.25 μM depth resolution.

To compare hemibrain neuron morphology to R65C03-GAL4 (Ext. Data Fig. 2c), we warped a stain of R65C03 (from Flylight^78^) and hemibrain skeletons of layer 6 tangential neurons using the 2018 Janelia female template brain^79^. We overlayed the warped stain and warped skeletons and identified neurons with high overlap.

### Data analysis

#### Connectomic analysis

Data from the hemibrain connectome^16^ were obtained from neuprint explorer (http://neuprint.janelia.org, hemibrain:v1.2.1) and analyzed using custom Python code. Prior to visualizing the weight matrices (Ext. Data Fig. 1f; Fig 5a), we sorted each cell type by their FB or protocerebral bridge x position (i.e. columns), with hΔK positioning determined by their axons. To count the fraction of FB6M or FB5V synapses onto hΔK axons, we sorted synapses by their x-position on a hΔK cell-by-cell basis. To calculate the ratio of recurrent synapse to EPG synapses (Ext. Data Fig. 9b), we also examined data from the FAFB (https://codex.flywire.ai, FAFB v783) and male CNS (http://neuprint.janelia.org, male-cns:v0.9) connectomes. For each PFG neuron, we calculated the total number of synapses from hΔK neurons, the total number of synapses to hΔK neurons, and the total number of synapses from EPG neurons. The recurrence was the average synapses to and from hΔK neurons, and this was divided by the total inputs from EPGs to calculate the ratio. Since the FAFB data was less curated, and occasionally contained PFG neurons with 0 connectivity to EPG or hΔK neurons, we only included PFG cells that had a total recurrence strength and heading strength both above 10.

#### Calcium imaging and behavior analysis

We used fictrac software^77^ to transform the x, y, and z rotations of a custom-made ball into 2D walking trajectories. The forward velocity and angular velocity were directly calculated as rotations around the x and z axes, respectively, which were then used to calculate heading (motor position, calculated as the integral of angular velocity), position (x = integral of forward velocity * sin(heading); y = integral of forward velocity * cos(heading)), and upwind velocity (vy = forward velocity * cos(heading)). The rotation around the y axis (side velocity) was additionally used to calculate travel direction (heading + tan^-1^(forward velocity / side velocity)). For data analysis, the position, angular velocity, angular speed, upwind velocity, and forward velocity were smoothed using a running average of 1 s width. Flies were only included for further analysis if they reliably oriented upwind on at least 50% of trials, which was identified by a convergence of heading values towards the upwind direction (0°) and an increase in mean upwind velocity around wind or odor onset.

We used custom Matlab software to pre-process our calcium imaging data (see ref^26^) for details), including motion correction, fluorescence extraction (across eight hand-drawn ROIs across the FB), and alignment with fictrac behavioral data. To detect activity bumps in hΔK neurons, PFG neurons, and neurons labeled by VT062617-Gal4, we low-pass filtered bump amplitude data (butterworth filter (fs = 100 Hz, fc = 2 Hz)) and then established a threshold on a trial-by-trial basis according to the minimum ΔF/F value of that trial and the ΔF/F standard deviation across all trials. For most analyses, this threshold was set at minimum(ΔF/F) + 1* standard deviation, but to identify large PFG bump onsets and offsets for Fig. 1k-m, the threshold was increased to minimum(ΔF/F) + 1.75 * standard deviation. A minimum bump duration was set at 3 s, and any above-threshold activity within 3 s of bumps was merged. To calculate bump position, the raw ΔF/F values across all eight ROIs were smoothed (running average, width ∼2 s) and then fit with a von Mises distribution at each time step using the Matlab function fminsearch. To aid fitting, we used the previous bump position or the ROI with the maximum ΔF/F (if the previous time step did not have a bump) as the initial search values. These bump positions were reported as a phase angle, where -180° is the left side of the FB and 180° is the right side of the FB.

During calcium imaging of FB5V and FB6M neurons, the calcium signal showed an artifactual decay from trial start to trial end. This artifact was removed by fitting an exponential decay using the beginning (first 5 s) and end (last 5 s) of the ΔF/F signal, and then subtracting this fit from the ΔF/F signal. Therefore, all ΔF/F signals reported for FB5V and FB6M are changes from this baseline.

In panels Fig. 1i,l, and Ext. Data Fig. 1k we calculated the persistence of walking direction following odor. To calculate the persistence of walking direction, we first defined a goal direction as the mean heading direction adopted in the first second after odor offset (post odor) or after trial start (baseline control), and then identified the time when flies stopped walking (forward velocity under 0.5 mm/s for at least 1.5 s) or deviated from the goal direction by at least 45°. In panels Fig 1h,i,k,l, we calculated the persistence of neural activity following odor. To calculate the neural persistence, we calculated the time between odor offset and bump offset. To better compare neural and behavioral persistence, we only considered trials where the bump activity turned on before odor offset and turned off after odor offset.

In panels Fig. 1j,m, we aligned bump amplitude and behavioral data to bump offset. We only included trials where the bump activity turned on before odor offset and turned off after odor offset, and we removed any data before bump onset. In control bump ON data, we took the same trials, but only included data between bump onset and bump offset, each aligned by the midpoint of the bump. For goal deviation, the ‘goal’ was defined as the mean heading during the first 1/6^th^ (between 0.18 – 4.46 s) of the bump ON or bump offset data.

In panel Fig 3d, we calculated the mean bump width of hΔK activity. To do this, we isolated periods with bumps, aligned the raw ΔF/F calcium signals (across 8 ROIs) to the column with the highest signal, and then normalized the activity by the maximum activity.

In panel Fig. 4d, we calculated the maximum bump deviation and cumulative bump deviation. To quantify maximum bump deviation, we calculated the difference between the minimum and maximum unwrapped bump position between each bump onset and offset. To quantify cumulative bump deviation, we calculated how much the unwrapped bump position changed at each time point between bump onset and offset, summed the absolute value of these changes, and then divided the sum by the total length of the bump period.

In panels Fig. 4e-g and Fig. 6f,g,o,p, we identified discrete periods of turning. To identify turns, we identified segments of data where the unwrapped heading shifted by at least 90°. We then identified the beginning and end of each turn by searching backwards or forwards in time, respectively, until the heading stabilized (the maximum change in heading over 0.25 seconds was smaller than the 80^th^ percentile of heading standard deviations across all bins of the same duration). Since this algorithm isolated periods of turning with different durations, when turns were aligned in time (Fig. 4g, Fig. 6f,o), the data were time warped such that all turns were equal in length. In cases where small turns were compared to large turns (Fig. 4g), turns were classified as small if the total change in heading was less than 130°, and large otherwise.

In panels Fig. 4e,f we further classified behavior as ‘rest’ or ‘straight running’. Rests were defined as periods of low movement (v < 0.5 mm/s) for at least 3 seconds. Straight running was defined as movements with high walking speed (v > 50th quartile of speed for given fly) and stable heading (Δheading < 90°). Short straight running bouts (under 3 s) within 1 s of larger bouts were merged, and large bouts (over 3 s) within 3 s of each other were merged. Any turns that had overlapping sections with straight running bouts were removed.

In Fig. 4h, we calculated discrete shifts in hΔK bump position following turns. To do this, we identified pairs of hΔK bumps separated in time. For each bump pair, we calculated the change in wrapped heading from the end of the first bump to the beginning of the second bump, and then classified the pairs as left turns (0° > Δ heading) or right turns (0° < Δ heading). Since these data were wrapped, we centered the wrapping around the bump position or heading taken at the end of the first bump.

In Fig. 4i-k, we calculated the rotational gain of the PFG bump as the cumulative change in unwrapped bump position divided by the cumulative change in unwrapped heading. As both bump position and heading are represented by angles (see above for bump position), a rotation gain of 0 indicates stable bumps and a rotation gain of 1 indicates perfect tracking. For Fig. 4i,j, this gain was calculated over entire bumps present during wind shifts. The average bump amplitude was calculated across this same time period. For Fig. 4k, rotational gain was calculated over a 1 s sliding window starting 5 seconds before odor onset and ending at odor offset. To avoid timepoints where the bump was mostly absent or the fly heading was stationary, we excluded timepoints if the total window did not contain sufficient bump data (< 50%), if the cumulative delta heading was too low (< 0.2), or if the rotational gain was too high (> 5).

In panels Ext. Data Fig. 8b-f, we correlated bump position with heading or travel direction for each trial. We only excluded trials if the trial contained no bump activity.

In panels Fig. 6e,n, we cross correlated FB5V and FB6M ΔF/F signals with angular speed or upwind velocity. Prior to cross correlating, we mean subtracted the fluorescent and behavioral signals. To remove wind-coupled activity fluctuations in FB6M data, we only used data from 5 s after wind onset and 5 s before wind offset.

In panels Fig. 6h,q, we aligned FB6M and FB5V activity to deviations from goals. We first defined a goal direction as the mean heading adopted in the first second after odor offset, and then aligned neural and behavioral data to the time when flies deviated from this goal by at least 45°.

#### Electrophysiology analysis

For analysis of electrophysiology data, the voltage traces were low pass filtered using a butterworth filter (fs = 10000 Hz, fc = 5 Hz), downsampled by a factor of 10, and then baseline subtracted. We calculated persistent excitation (Fig. 2d,f,i; Ext. Data Fig. 3e,h) by summing the baseline-subtracted voltage signal between light offset and trial end, and then normalizing by the length of this period. For visualizing voltage signals across trials (e.g., Fig. 2c), the voltage signals were sorted by persistent excitation (Fig. 2a,c,e,g,h; Ext. Data Fig. 3a) or inhibition during the stimulus (Fig. 2j,l; Ext. Data Fig. 4c).

#### Statistics

All statistics were ran using GraphPad Prism 10 and reported in Tables 3 and 4. Non-parametric statistical tests were used if the data was not normally distributed (see Tables 3 and 4). An alpha value of 0.05 was used to determine significance, with *0.01<p<0.05, **0.001<p<0.01, ***0.0001<p<0.001, ****p<0.0001 reported on figures.

**Table 3.** Drug sources and concentrations.

| Drug | Final concentration | Source | Identifier |
| --- | --- | --- | --- |
| Tubocurarine chloride (curare) | 10 μM | Tocris | 2820 |
| Imidacloprid (IMI) | 100 nM | Sigma | 37894 |
| Picrotoxin | 5 μM | Tocris | 1128 |
| Methysergide | 50 μM | Sigma | M137 |

**Table 4:** Statistics for main figures. Significant p values are bolded.

| **Figure** | **Comparison** | **Sample size** | **Statistical test** | **p value** |
| --- | --- | --- | --- | --- |
| Fig. 1f | base vs wind | N=6 flies<br>n = 39-49 trials/fly | Paired t-test | **0.0027** |
|  | base vs odor | N=6 flies<br>n = 39-49 trials/fly | Paired t-test | **0.0006** |
| Fig. 1g | base vs wind | N=12 flies<br>n = 36-60 trials/fly | Paired t-test | **0.0021** |
|  | base vs odor | N=12 flies<br>n = 36-60 trials/fly | Paired t-test | **0.0003** |
| Fig. 1h | Average bump amplitude | N = 6 flies<br>n = 39-49 trials/fly | N/A | N/A |
| Fig. 1i | persistence distributions | N = 6 flies<br>n = 18-29 trials/fly | N/A | N/A |
| Fig. 1j | goal deviation bump vs offset | N = 6 flies<br>n = 12-23 trials/fly | Paired t-test | **0.0017** |
|  | forward velocity compared to 0 | N = 6 flies<br>n = 12-23 trials/fly | Paired t-test | 0.0522 |
|  | P(upwind heading) compared to 0 | N = 6 flies<br>n = 12-23 trials/fly | Paired t-test | 0.0694 |
| Fig. 1k | Average bump amplitude | N = 6 flies<br>n = 49-59 trials/fly | N/A | N/A |
| Fig. 1l | persistence distributions | N = 12 flies<br>n = 8-41 trials/fly | N/A | N/A |
| Fig. 1m | goal deviation bump vs offset | N = 12 flies<br>n = 2-31 trials/fly | Paired t-test | **0.0028** |
|  | forward velocity compared to 0 | N = 12 flies<br>n = 2-31 trials/fly | Paired t-test | **0.0295** |
|  | P(upwind heading) compared to 0 | N = 12 flies<br>n = 2-31 trials/fly | Paired t-test | 0.1229 |
| Fig. 2d | pre-drug vs IMI + curare | N = 8 cells<br>n<sub>pre-drug</sub> = 6-10 trials/cell<br>n<sub>IMI_curare</sub> = 1-16 trials/cell | Wilcoxon matched-pairs signed rank test | **0.0391** |
| Fig. 2f | control vs hΔK>TNTe | N<sub>control</sub> = 15 cells<br>n<sub>control</sub> = 5-10 trials/cell<br>N<sub>TNTe</sub> = 8 cells<br>n<sub>TNTe</sub> = 1-9 trials/cell | Mann Whitney signed-rank test | **0.0032** |
| Fig. 2i | pre-drug vs methysergide | N = 8 cells<br>n<sub>pre-drug</sub> = 5-8 trials/cell<br>n<sub>methysergide</sub> = 5-9 trials/cell | Paired t-test | **0.0367** |
| Fig. 2k | pre-drug vs picrotoxin | N = 5 cells<br>n<sub>pre-drug</sub> = 5-10 trials/cell<br>n<sub>picrotoxin</sub> = 2-6 trials/cell | Paired t-test | **0.0039** |
| Fig. 2m | pre-drug vs picrotoxin | N = 4 cells<br>n<sub>pre-drug</sub> = 4-7 trials/cell<br>n<sub>picrotoxin</sub> = 2-9 trials/cell | Paired t-test | **0.0212** |
| Fig. 3d | Columns vs 0 | N = 6 flies<br>n = 39-49 trials/fly | One-way ANOVA<br>with multiple<br>comparisons | **Main effect of column:**<br>$p < 0.0001$<br>**Column 4 vs -3:**<br>$p = 0.9410$<br>**Column 4 vs -2:**<br>$p = 0.7244$<br>**Column 4 vs -1:**<br>$p = 0.0006$<br>**Column 4 vs -0:**<br>$p < 0.0001$<br>**Column 4 vs 1:**<br>$p = 0.0013$<br>**Column 4 vs 2:**<br>$p = 0.9018$<br>**Column 4 vs 3:**<br>$p = 0.9588$ |
| Fig. 4d | Max bump deviation hΔK vs PFGs | N<sub>hΔK</sub> = 281 bumps<br>N<sub>PFG</sub> = 695 bumps | Mann Whitney<br>signed-rank test | $< 0.0001$ |
| | Cumulative bump deviation hΔK vs PFGs | N<sub>hΔK</sub> = 281 bumps<br>N<sub>PFG</sub> = 695 bumps | Mann Whitney<br>signed-rank test | $< 0.0001$ |
| Fig. 4e | hΔK vs PFGs | hΔK:<br>N = 6 flies<br>n<sub>rest</sub> = 6-73<br>rests/fly<br>n<sub>turn</sub> = 57-76<br>turns/fly<br>n<sub>straight_run</sub> = 2-93<br>runs/fly<br>PFGs:<br>N = 12 flies<br>n<sub>rest</sub> = 19-87<br>rests/fly<br>n<sub>turn</sub> = 24-181<br>turns/fly<br>n<sub>straight_run</sub> = 7-75<br>runs/fly | Two-way analysis<br>of variance with<br>multiple<br>comparisons | **Main effect of behavioral stage:**<br>$p < 0.0001$<br>**Main effect of genotype:**<br>$p < 0.0001$<br>**Interaction:**<br>$p = 0.0470$<br>**Rest hΔK vs PFGs:**<br>$p < 0.0001$<br>**Turn hΔK vs PFGs:**<br>$p < 0.0001$<br>Surge hΔK vs PFGs:<br>$p = 0.1549$ |
| Fig. 4f | hΔK across behavioral stage | N = 6 flies<br>n<sub>rest</sub> = 6-73<br>rests/fly<br>n<sub>turn</sub> = 57-76<br>turns/fly<br>n<sub>straight_run</sub> = 2-93 | Friedman test with<br>multiple<br>comparisons | **Main effect of behavioral stage:**<br>$p = 0.0081$<br>rest vs turn:<br>$p > 0.9999$ |
| | | runs/fly | | **rest vs straight running:**<br>$p = 0.0281$<br>**turn vs straight running:**<br>$p = 0.0281$ |
| | PFGs across behavioral stage | N = 12 flies<br>$n_{\text{rest}} = 19-87$ rests/fly<br>$n_{\text{turn}} = 24-181$ turns/fly<br>$n_{\text{straight\_run}} = 7-75$ runs/fly | One-way analysis of variance with multiple comparisons | **Main effect of behavioral stage:**<br>$p < 0.0001$<br>rest vs turn:<br>$p = 0.4704$<br>**rest vs straight running:**<br>$p < 0.0001$<br>**turn vs straight running:**<br>$p < 0.0001$ |
| Fig. 4g | PFG across turns | Left turn ($>130^\circ$)<br>N = 139 turns<br>Left turn ($<130^\circ$)<br>N = 111 turns<br>Right turn ($<130^\circ$)<br>N = 198 turns<br>Right turn ($>130^\circ$)<br>N = 177 turns | Wilcoxon signed rank test | **Left turn ($>130^\circ$):**<br>$p < 0.0001$<br>**Left turn ($<130^\circ$):**<br>$p < 0.0001$<br>**Right turn ($<130^\circ$):**<br>$p < 0.0001$<br>**Right turn ($>130^\circ$):**<br>$p < 0.0001$ |
| Fig. 4h | hΔK across turns | Left turns:<br>N = 476 turns<br>Right turns:<br>N = 696 turns | circ_medtest from Circular Statistics Toolbox (test significance from median = 0) | **Left turns:**<br>$p = 0.0089$<br>**Right turns:**<br>$p < 0.0001$ |
| Fig. 4j | rotation gain vs bump amplitude | N = 83 bumps | correlation | **$r = -0.4032$**<br>**$p = 0.0002$** |
| Fig. 4k | base vs odor | N = 12 flies<br>$n = 36-60$ trials/fly | Paired t-test | **$p = 0.0013$** |
| Fig. 6c | mean $\Delta F/F$ | N = 6 flies<br>$n = 8-13$ trials/fly | N/A | N/A |
| Fig. 6e | angular speed cross correlation | N = 6 flies<br>$n = 39-65$ trials/fly | One-sample t-test | **0.0019** |
| | upwind velocity cross correlation | N = 6 flies<br>$n = 39-65$ trials/fly | One-sample t-test | **0.0081** |
| Fig. 6f | mean \|heading\| and $\Delta F/F$ | N = 322 turns | N/A | N/A |
| Fig. 6g | base vs turn | N = 322 turns | Wilcoxon matched-pairs signed rank test | **$< 0.0001$** |
| Fig. 6h | mean goal | N = 6 flies | N/A | N/A |
| | deviation and $\Delta F/F$ | n = 39-65 trials/fly | | |
| Fig. 6i | base vs goal offset | N = 6 flies<br>n = 39-65 trials/fly | Paired t-test | **0.0423** |
| Fig. 6l | mean $\Delta F/F$ | N = 6 flies<br>n = 4-12 trials/fly | N/A | N/A |
| Fig. 6n | angular speed<br>cross correlation | N = 6 flies<br>n = 26-56 trials/fly | One-sample t-test | **0.0039** |
|  | upwind velocity<br>cross correlation | N = 6 flies<br>n = 26-56 trials/fly | One-sample t-test | 0.4382 |
| Fig. 6o | mean \|heading\|<br>and $\Delta F/F$ | N = 322 turns | N/A | N/A |
| Fig. 6p | base vs turn | N = 180 turns | Wilcoxon matched-pairs signed rank<br>test | **< 0.0001** |
| Fig. 6q | mean goal<br>deviation and $\Delta F/F$ | N = 6 flies<br>n = 26-56 trials/fly | N/A | N/A |
| Fig. 6r | base vs goal offset | N = 6 flies<br>n = 26-56 trials/fly | Wilcoxon matched-pairs signed rank<br>test | **0.0312** |

**Table 5:** Statistics for Extended Data figures. Significant p values are bolded.

| Figure | Comparison | Sample size | Statistical test | p value |
| --- | --- | --- | --- | --- |
| Ext. Data Fig. 1j | heading distributions | N = 18 flies<br>n<sub>baseline</sub> = 935 trials<br>n<sub>odor</sub> = 935 trials<br>n<sub>10s</sub> = 935 trials<br>n<sub>20s</sub> = 935 trials<br>n<sub>30s</sub> = 724 trials | N/A | N/A |
| Ext. Data Fig. 1k | trajectory persistence distributions | N = 18 flies<br>n = 10-48 trials/fly | N/A | N/A |
| Ext. Data Fig. 2b | mean upwind velocity and bump amplitude | N = 6 flies<br>n<sub>1</sub> = 3-7 trials/fly<br>n<sub>2</sub> = 2-8 trials/fly<br>n<sub>3</sub> = 2-9 trials/fly<br>n<sub>4</sub> = 4-8 trials/fly | N/A | N/A |
| Ext. Data Fig. 2c | mean upwind velocity and bump amplitude | N = 6 flies<br>n<sub>1</sub> = 6-12 trials/fly<br>n<sub>2</sub> = 5-12 trials/fly<br>n<sub>3</sub> = 7-12 trials/fly<br>n<sub>4</sub> = 5-12 trials/fly | N/A | N/A |
| Ext. Data 2d | Mean bump amplitude before and after odor | N<sub>hΔK</sub> = 6 flies<br>n<sub>hΔK</sub> = 39-49 trials/fly<br>N<sub>PFG</sub> = 12 flies<br>n<sub>PFG</sub> = 36-60 trials/fly | Paired t-test | **p<sub>hΔK</sub> = 0.0397**<br>**p<sub>PFG</sub> < 0.0001** |
| Ext. Data Fig. 3b | mean Δvoltage across current steps | N = 32 cells<br>n<sub>-4pA</sub> = 3-10 trials/cell<br>n<sub>0pA</sub> = 3-10 trials/cell<br>n<sub>4pA</sub> = 4-9 trials/cell | N/A | N/A |
| Ext. Data Fig. 3d | mean voltage over time | N = 13 cells<br>n<sub>0.2</sub> = 4-9 trials/cell<br>n<sub>0.8</sub> = 5-9 trials/cell | N/A | N/A |
| Ext. Data Fig. 3e | light duty cycle | N = 13 cells<br>n = 4-9 trials/cell<br>n<sub>0.5</sub> = 4-9 trials/cell<br>n<sub>0.8</sub> = 5-9 trials/cell | Friedman test with multiple comparisons | **Main effect of duty cycle:**<br>**p < 0.0001**<br>**0.2 vs 0.5:**<br>**p = 0.0181**<br>**0.2 vs 0.8:**<br>**p < 0.0001**<br>0.5 vs 0.8:<br>p = 0.3500 |
| Ext. Data Fig. 3f | mean voltage over time | N = 8 cells<br>n<sub>0.2</sub> = 1-7 trials/cell<br>n<sub>0.8</sub> = 2-7 trials/cell | N/A | N/A |
| Ext. Data Fig. 3g | mean voltage over time | N = 13 cells<br>n<sub>0.2</sub> = 5-8 trials/cell<br>n<sub>0.8</sub> = 5-9 trials/cell | N/A | N/A |
| Ext. Data Fig. 3h | light duty cycle | N = 13 cells<br>n<sub>0.2</sub> = 5-8 trials/cell<br>n<sub>0.5</sub> = 4-8 trials/cell<br>n<sub>0.8</sub> = 5-9 trials/cell | Friedman test with multiple comparisons | **Main effect of duty cycle:**<br>**p = 0.0014**<br>**0.2 vs 0.5:** |
|  |  |  |  | **$p = 0.0239$**<br>**0.2 vs 0.8:**<br>**$p = 0.0016$**<br>0.5 vs 0.8:<br>$p > 0.9999$ |
| Ext. Data<br>Fig. 3i | mean voltage<br>over time | N = 8 cells<br>$n_{0.2} = 3-5$ trials/cell<br>$n_{0.8} = 3-5$ trials/cell | N/A | N/A |
| Ext. Data<br>Fig. 4d | pre-drug vs<br>picrotoxin | N = 6 cells<br>$n_{\text{pre-drug}} = 5-7$ trials/cell<br>$n_{\text{picrotoxin}} = 3-13$ trials/cell | Paired t-test | **0.0049** |
| Ext. Data<br>Fig. 8d | $h\Delta K$ and PFGs<br>vs 0 | $N_{h\Delta K} = 231$ trials<br>$N_{\text{PFGs}} = 447$ trials | Wilcoxon<br>signed rank<br>test | **$h\Delta K$ vs 0:**<br>**$p < 0.0001$**<br>**PFGs vs 0:**<br>**$p < 0.0001$** |
| Ext. Data<br>Fig. 8e | $h\Delta K$ vs PFGs | $N_{h\Delta K} = 231$ trials<br>$N_{\text{PFGs}} = 447$ trials | Wilcoxon<br>signed rank<br>test | $h\Delta K$ vs 0:<br>$p = 0.2166$<br>**PFGs vs 0:**<br>**$p = 0.0002$** |
| Ext. Data<br>Fig. 8f | $h\Delta K$ heading<br>vs travel<br>direction | N = 231 trials | Wilcoxon<br>matched-pairs signed<br>rank test | **$p = 0.0141$** |
|  | PFGs heading<br>vs travel<br>direction | N = 447 trials | Wilcoxon<br>matched-pairs signed<br>rank test | **$p < 0.0001$** |
| Ext. Data<br>Fig. 10a | mean $\Delta F/F$<br>over time | N = 6 flies<br>$n_1 = 8-14$ trials/fly<br>$n_2 = 7-12$ trials/fly<br>$n_3 = 8-14$ trials/fly<br>$n_4 = 8-12$ trials/fly | N/A | N/A |
| Ext. Data<br>Fig. 10b | mean $\Delta F/F$<br>over time | N = 6 flies<br>$n_1 = 5-11$ trials/fly<br>$n_2 = 5-11$ trials/fly<br>$n_3 = 5-11$ trials/fly<br>$n_4 = 5-11$ trials/fly | N/A | N/A |

### Modeling

#### Full FB model

To model the FB recurrent network, we created a rate-based dynamical system model consisting of 30 hΔK neurons, 18 PFG neurons, and 1 global inhibitory neuron. We approximated all dynamical systems using Euler’s method of integration with a time step of 0.001 s. To generate ring recurrence between hΔK and PFG neurons, we gave each neuron a relative position in the FB, indicated by an angle from 0 to 2π, and then connected the populations with local excitation that was maximum at overlapping positions and decayed with distance. This was done explicitly using a Gaussian equation fit to the real connectivity from the hemibrain connectome (Fig. 3; Fig. 5) or using the connectivity directly from the connectome (Ext. Data Fig. 6). The local spread of the Gaussian connectivity was set by the local connectivity width (σ, Gaussian standard deviation), which establishes how quickly the connectivity decays as the distance (in radians) between hΔK and PFG neurons increases (σ_PFG > hΔK_ = 0.25, σ_hΔK > PFG_ = 0.25, in radians). The best fit Gaussian connectivity was found by searching for the local connectivity width (σ) that minimized the absolute error between the fit weight matrix and real weight matrix. Each column of the real weight matrix (the connectivity of each PFG neuron to all hΔK neurons) was normalized to ensure fits were based on the local connectivity width rather than variability in synaptic strength across the FB. Although real hΔK neurons have a 180° shift across the FB, for simplicity, we only show connectivity and activity simulations from the ‘axons’ of hΔK neurons. The weight matrices between hΔK or PFGs and the global inhibitory neuron were either one-to-all or all-to-one. To simulate the neural activity of each population, we used the following ordinary differential equations (ODEs):

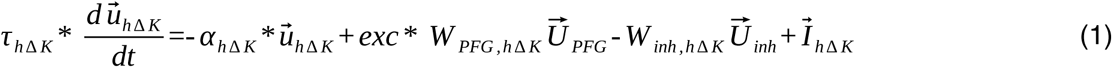

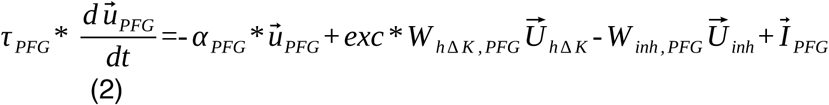

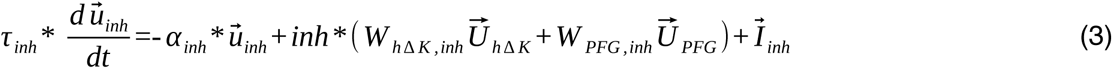

Where *τ_x_* is the time constant for population x, *α_x_* is the leak of population x, *u⃗*_*x*_ is the activity of population x, *exc* is the excitation strength, *inh* is the inhibition strength, *W_x,y_* is the weight matrix from population x to population y, *U⃗*_*x*_ is the synaptic output of population x, and *I⃗*_*x*_ is the external input to population x. *τ_x_* and *α_x_* were set to 1 and 10, respectively, for all populations. *I⃗*_*x*_ was used to set the input patterns to each population.

To simulate the synaptic output of each population, we used the following second order ODEs to simulate alpha functions:

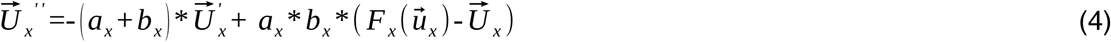

Where *1/a_x_* is the decay rate of the alpha function for population x, *1/b_x_* is the growth rate of the alpha function for population x, and *F_x_* is the activation function of population x. The synaptic growth and decay values used for fast and slow synapses were fit to synaptic physiology data by searching for the *a* and *b* values that minimized the absolute error between the fit activity and synaptic physiology voltage data. The fast synapses were fit to hΔK voltage responses to FB6M (49B08AD; VT063626DBD) pulsed stimulation, and were set to *a_x_* = 4.63 and *b_x_* = 25. The slow synapses were fit to hΔK voltage responses to FB6A/D (65C03-Gal4) stimulation in the presence of curare (10 µM) and IMI (100 nM), and were set to *a_x_* = 0.64 and *b_x_* = 25. We used the following activation functions:

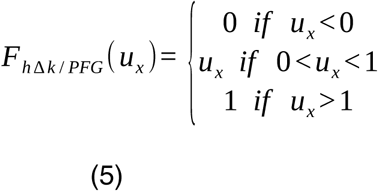

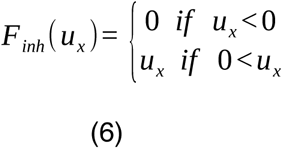

During the simulations in Fig. 3, sinusoidal (direct offset = 1), Gaussian (normalized such that max = 1), or random input (Gaussian distribution with µ = 1, σ = 0.5) were provided to the hΔK and PFG neurons for 4 or 10 seconds. During the simulations in Ext. Data Fig. 8a, Fig. 5, and Ext. Data Fig. 9e we only provided input to PFG neurons, which was sinusoidal with a phase that we randomly and stochastically updated over time. Phase shifts were stochastic, occurring with a 0.1% chance at each time step, and when they occurred, they were randomly drawn from a gaussian distribution with µ = 0 and σ = 50° and then smoothed over time according to:

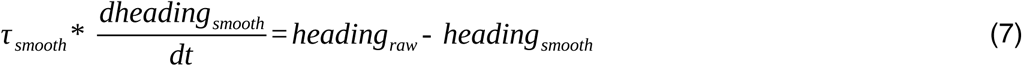

Where *τ_smooth_* equals 1. To account for the differences in synaptic strength between EPGs > PFG and hΔK <-> PFG synapses, we decreased the amplitude of the sinusoidal input to 1/6 of the recurrent excitation strength. We controlled the gate in Fig. 5 by either externally changing the excitation strength (*exc* in equations 1,2, increased from 0 to 7) or changing external input to hΔK (*I⃗*_*hΔK*_ in equation 1, inhibition corresponded with a value of -5, and disinhibition corresponded with a value of 0). In Ext. Data Fig. 9e, we additionally attempted to gate bump dynamics by adding external input to hΔK and PFG neurons (*I⃗*_*hΔK*_ and *I⃗*_*PFG*_ in equation 1, increased from -1 or -2 to 0).

To calculate the network decay rate in Fig. 3, we calculated the maximum difference in hΔK output at each time step following external input offset and then fit this difference with an exponential decay function.

In Fig. 3d, we expanded the model simulations to predict how each simulation would appear if the neurons were recorded using population imaging, such as calcium imaging. Calcium imaging captures activity throughout each active neurons’ axons. Therefore, we calculated how much each model neuron should spread across the FB to synapse with its postsynaptic partners. This spread was then applied to the model simulation to get a better prediction of how far activity of the active neurons would spread across the FB. In Ext. Data Fig. 5f,g, to keep the neuron size the same across simulations we used real connectome data to estimate the width of real hΔK neurons, and then used this spread to estimate the width of FB activity. We fit the relationship between local connectivity standard deviation and FB activity percentage with an exponential equation.

In the parameter space in Ext. Data Fig. 7c,d, for each pair of parameters (a_inhibition_ and a_excitation_), we first calculated the decay tau of the excitatory neuron across each excitatory strength value, removed all values above 30 s, and then we summed these values to get the total ‘sum’.

In Fig. 5f and Ext. Data Fig. 6, we calculated the bump position by finding the relative position (as an angle) of the neuron with the maximum activity.

#### Simple recurrent network model

To simplify the recurrent network to two dynamic variables, we began by reducing the hΔK and PFG populations to one global excitatory neuron. This led to the following 6 variable dynamical system:

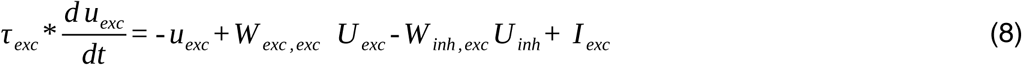

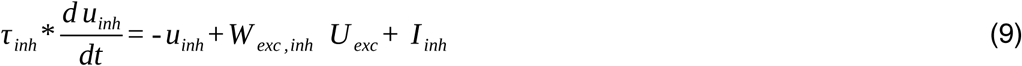

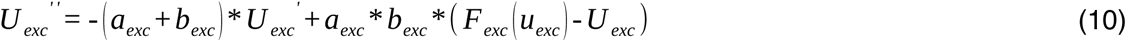

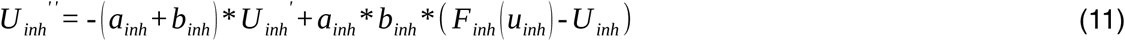

We made two simplifying assumptions. First, we assumed that the synaptic rise and decay constants (*1/a_x_, 1/b_x_*) were much slower than the membrane time constants (*τ_x_*). This means that the variables *u_exc_* and *u_inh_* instantaneously follow the synaptic dynamics. Second, we assumed that the synaptic decay time constant (*a_x_*) was much slower than the synaptic rise time constant (*b_x_*). These assumptions lead to the following 2 variable dynamical system:

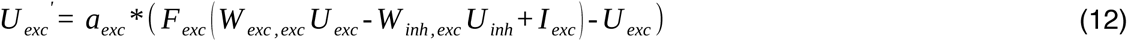

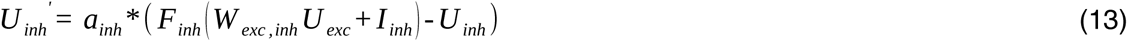

We used the same activation functions as those used for the excitatory and inhibitory neurons in the full model (equations 5,6). During simulations, we kept *W_exc,inh_* constant at 1, whereas we varied *W_exc,exc_* and *W_inh,exc_* with excitation strength and inhibition strength, respectively. During input, the excitatory unit received an external input equal to 1, whereas the inhibitory unit received no input.

## Data Availability Statement

All data will be made available on Zenodo upon publication.

## Code Availability Statement

All code will be made available at Github upon publication.

## Acknowledgements

The authors would like to thank Niels Ringstad, Dima Rinberg, Nicholas Tristch, XJ Wang for input on the project, and Michael Long, Jonathan Victor, Floris van Breugel, Mubarak Syed, and members of the Nagel and Schoppik labs for input on the manuscript. Kavin Nuñez warped stains of R65C03 and connectome skeletons. Alex Bates provided access to neurotransmitter predictions for hemibrain cell types prior to publication. Tanya Wolff provided split lines for hΔK prior to publication. Several attendees of the KITP workshop on Neurophysics of Active Sensing, including David Kleinfeld, provided helpful feedback on the Discussion. Eleni Samara provided help with Flywire images. This work was supported by NINDS Brain Initiative NS127129, NIDCD DC017979, and NSF 2014217 Odor2Action to K.I.N.

## Author Contributions

A.J.D. and K.I.N designed the study. A.J.D. performed all electrophysiology and imaging experiments, analyzed all data, and performed all computational modeling. N.D.K. provided assistance and training on the 2P imaging protocol and contributed to data analysis. E.H performed confocal imaging. B.E. provided advice and feedback on computational modeling and developed the minimal recurrent circuit model. A.J.D. and K.I.N. wrote the paper with feedback from all authors.

## Materials and Correspondence

Correspondence and requests for materials should be addressed to Katherine Nagel at.

## Extended Data

**Ext. Data Fig. 1:**
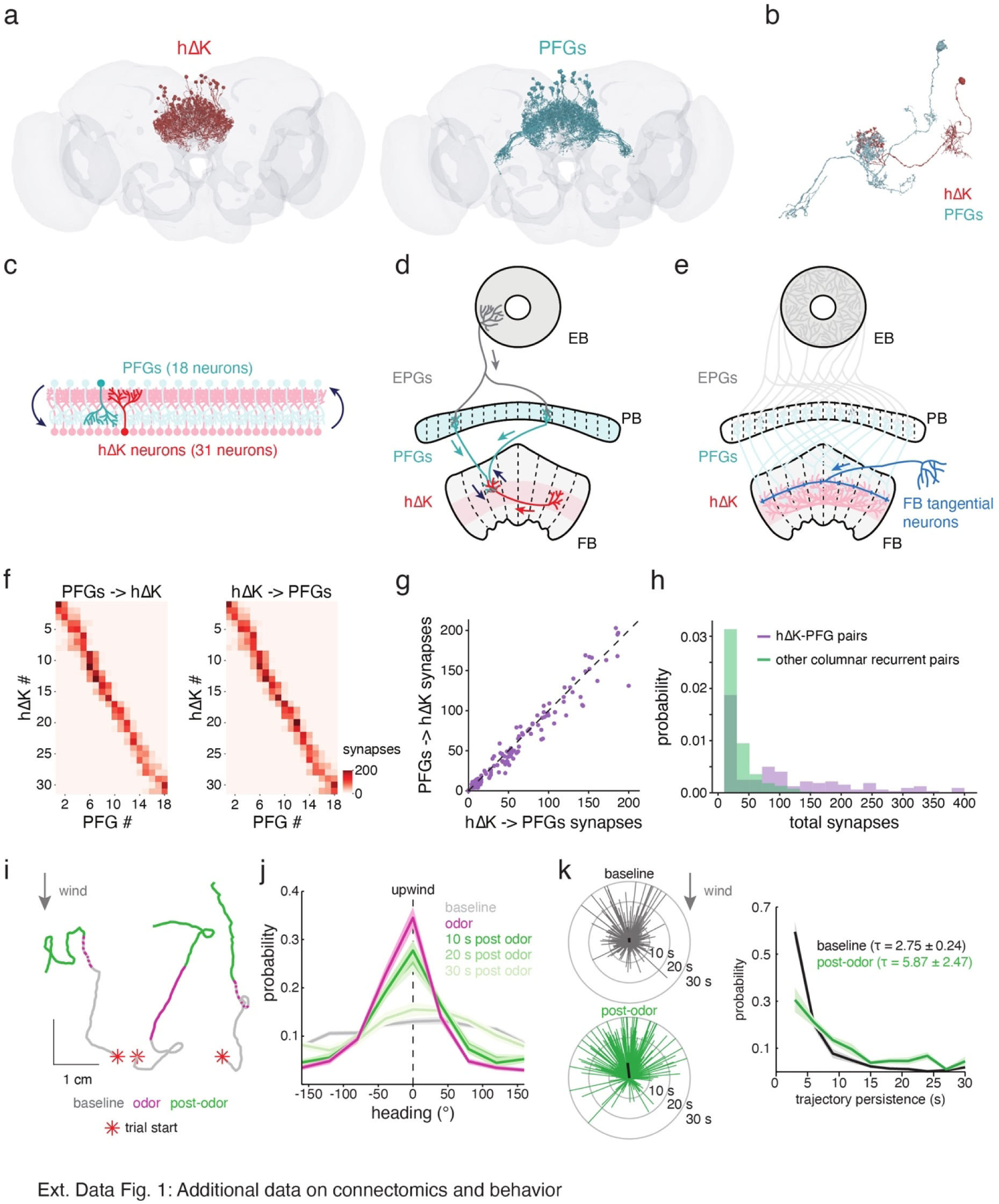
Additional data on connectomics and neural behavior. **a,** Morphology of the hΔK population (left) and PFGs population (right) from the FAFB connectome. **b,** Morphology of a connected pair of hΔK and PFG neurons. **c**, Schematic of the axo-axonic connectivity between hΔK (31 neurons) and PFGs (18 neurons). Each hΔK neuron synapse onto fewer PFG neurons than vice versa to account for the difference in population sizes. **d,** Schematic of the connectivity between EPG, PFG, and hΔK neurons. Individual EPG neurons synapse with two sets of PFG neurons in each half of the protocerebral bridge (PB). These PFG neurons reconverge back onto one part of the FB to reciprocally synapse with axons of hΔK neurons. Note, PFG neurons also receive heading inputs from inhibitory Δ7 neurons in the PB, with the same functional phase as EPG inputs (see Fig. 5 for connectivity). **e,** Schematic of the connectivity between FB tangential neurons and hΔK and PFG populations. FB tangential neurons send a single process into the FB that innervates all members of hΔK and/or PFG neurons. **f,** Synaptic weights from PFG to hΔK neurons (left) and hΔK to PFG neurons (right) taken from the hemibrain connectome. Both hΔK and PFG neurons are sorted by their axon position along the FB. **g,** Comparison of the reciprocal synaptic weights between hΔK and PFG neurons. Each dot represents a pair of hΔK and PFG neurons. **h,** Distributions of the total synaptic weight between pairs of recurrently-connected columnar neurons in the central complex. Data are separated for hΔK-PFG pairs (purple) and other columnar pairs (green), which includes all EPG, Δ7, PFN, PFL, PFR, FC, FS, FR, hΔ, and vΔ neurons. **i,** Example virtual trajectories of walking flies before (grey), during (pink), and following (green) odor. Note how flies often maintain their walking direction following odor offset (middle, right). **j,** Distribution of heading before wind/odor (baseline), during odor, 10 s after odor, 20 s after odor, and 30 s after odor (error = mean ± SEM), where 0° indicates upwind. Asterisks indicate significant difference from baseline distribution within shaded region. **k,** Left, vectors representing the direction (angle) and duration (length) flies walk following trial start (top, baseline) or odor offset (bottom), with upwards indicating upwind. The black lines represent the mean vector for each condition. Note, in the baseline condition, if flies maintained their direction for long enough following trial start, wind and odor were turned on, leading to the slight bias towards upwind in baseline, particularly for long walks. Right, distributions of the persistence of walking trajectories following trial start (baseline) or odor offset.

**Ext. Data Fig. 2:**
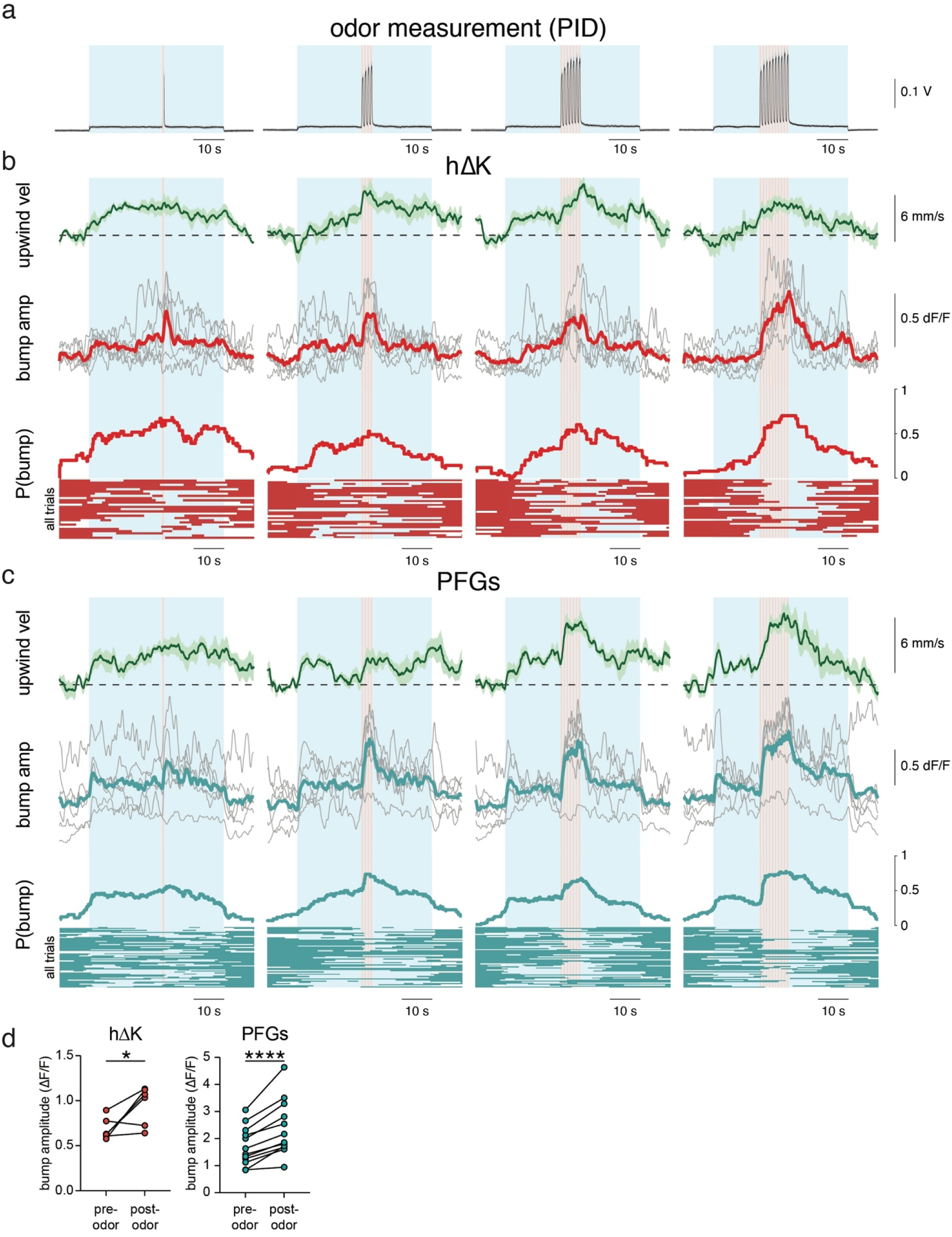
Additional data on neural responses of hΔK and PFGs. **a,** Photoionization detector (PID) measurements of odor concentration made at the location of the fly during baseline (no stimuli), wind, and pulses (1, 4, 7, or 10) of odor. **b,** Mean upwind velocity (walking aligned with wind), hΔK bump amplitude, and hΔK bump probability during baseline (no stimuli), wind, and pulses (1, 4, 7, or 10) of odor. Individual fly averages in grey; average across flies in green (behavior) and red (neural activity). Shaded regions are mean ± SEM. Below P(bump) is a binarized bump data for each trial, where red is no bump and transparent is bump. **c,** Same as **b** but for PFG neurons. Individual fly averages in grey; average across flies in green (behavior) and blue (neural activity). Below P(bump) is a binarized bump data for each trial, where blue is no bump and transparent is bump. **d**, Average bump amplitude in the 2.5 s before odor onset compared to in the 2.5 s following odor offset for hΔK (left) and PFG (right).

**Ext. Data Fig. 3:**
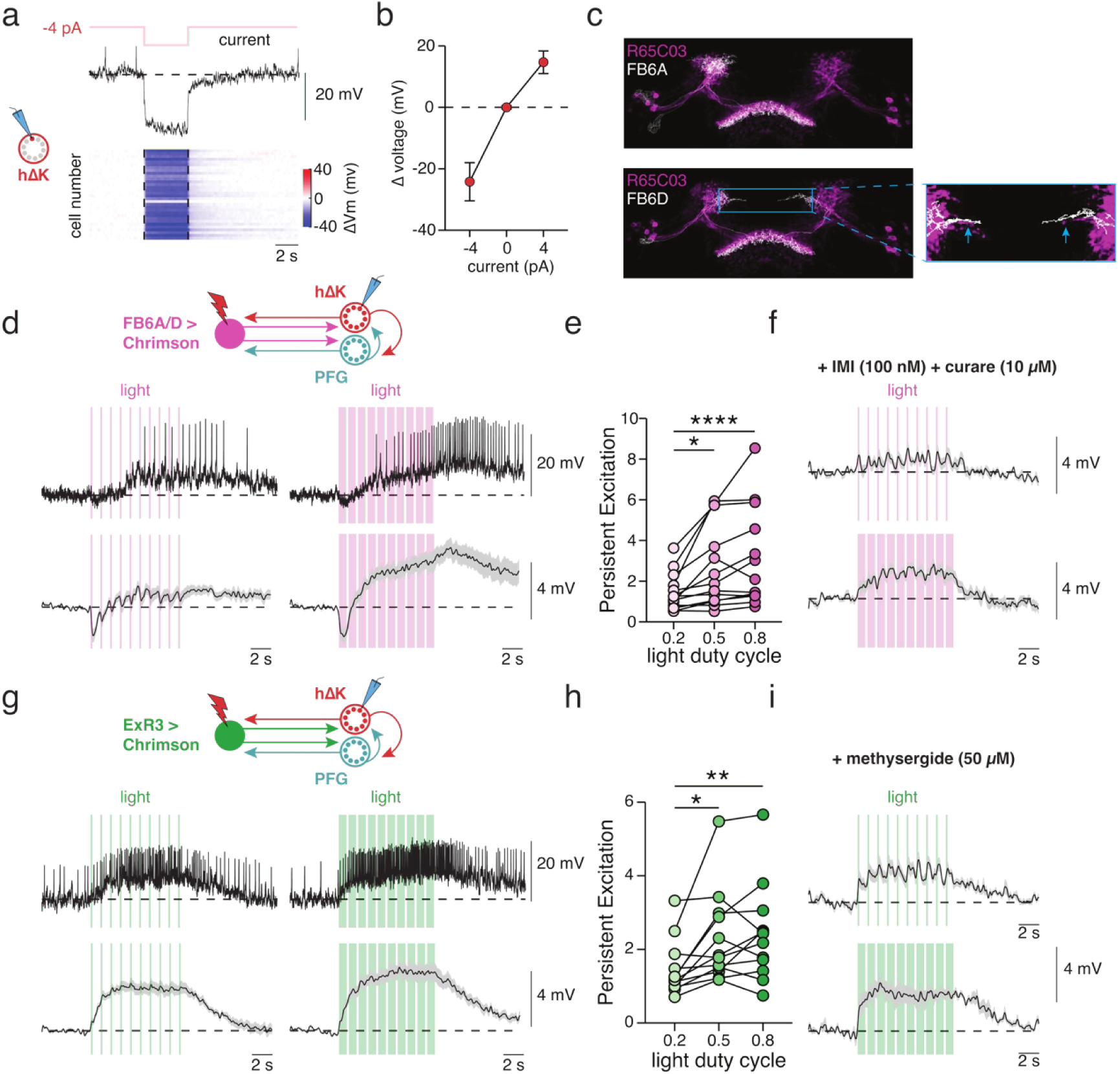
Additional data on excitatory inputs to hΔK. **a,** Voltage response of hΔK neurons (VT062617-lexA) during a -4 pA current injection. Top: example trace, bottom: heat map of mean membrane potential responses with each cell as a row. **b,** Mean membrane potential responses of hΔK neurons during -4pA, 0pA, or 4pA current injections. **c,** Overlay of R65C03 GAL4 line (image from Flylight^77^) and FB6A (top) or FB6D (bottom) skeletons from the hemibrain connectome, all warped to the Janelia female template brain^78^. The dendrites of FB6A and FB6D overlap strongly with the R65C03 stain. The right shows that R65C03 contains the medial-projecting processes that are unique to FB6D (blue arrows). Note that R65C03 contains multiple tangential neurons (around 17 per side), which overlaps with several other layer 6 neurons, such as FB6C_a, FB6C_b, FB6E, FB6G, FB6I, and FB6Z. **d,** Spiking and membrane potential responses of hΔK neurons to optogenetic activation of FB6A/D (65C03-gal4) using pulsed light (1 Hz) at 0.2 (left) or 0.8 (right) duty cycle. **e,** Mean persistent excitation (see Methods) following activation of FB6A/D at 0.2, 0.5, or 0.8 duty cycle. Data for 0.5 duty cycle are from Fig. 2c. **f,** Mean membrane potential responses of hΔK neurons in response to optogenetic activation of FB6A/D using pulsed light (1 Hz) at 0.2 (top) or 0.8 (bottom) duty cycle in the presence of IMI (100 nM) and curare (10 µM). **g,** Same as **d** but in response to activating ExR3 neurons (24C07-gal4). **h,** Same as **e**, but in response to activating ExR3 neurons. Data for 0.5 duty cycle are from Fig. 2g. **i,** Mean membrane potential of hΔK neurons in response to optogenetic activation of ExR3 using pulsed light (1 Hz) at 0.2 (top) or 0.8 (bottom) duty cycle in the presence of the general serotonin receptor antagonist methysergide (50 µM).

**Ext. Data Fig. 4:**
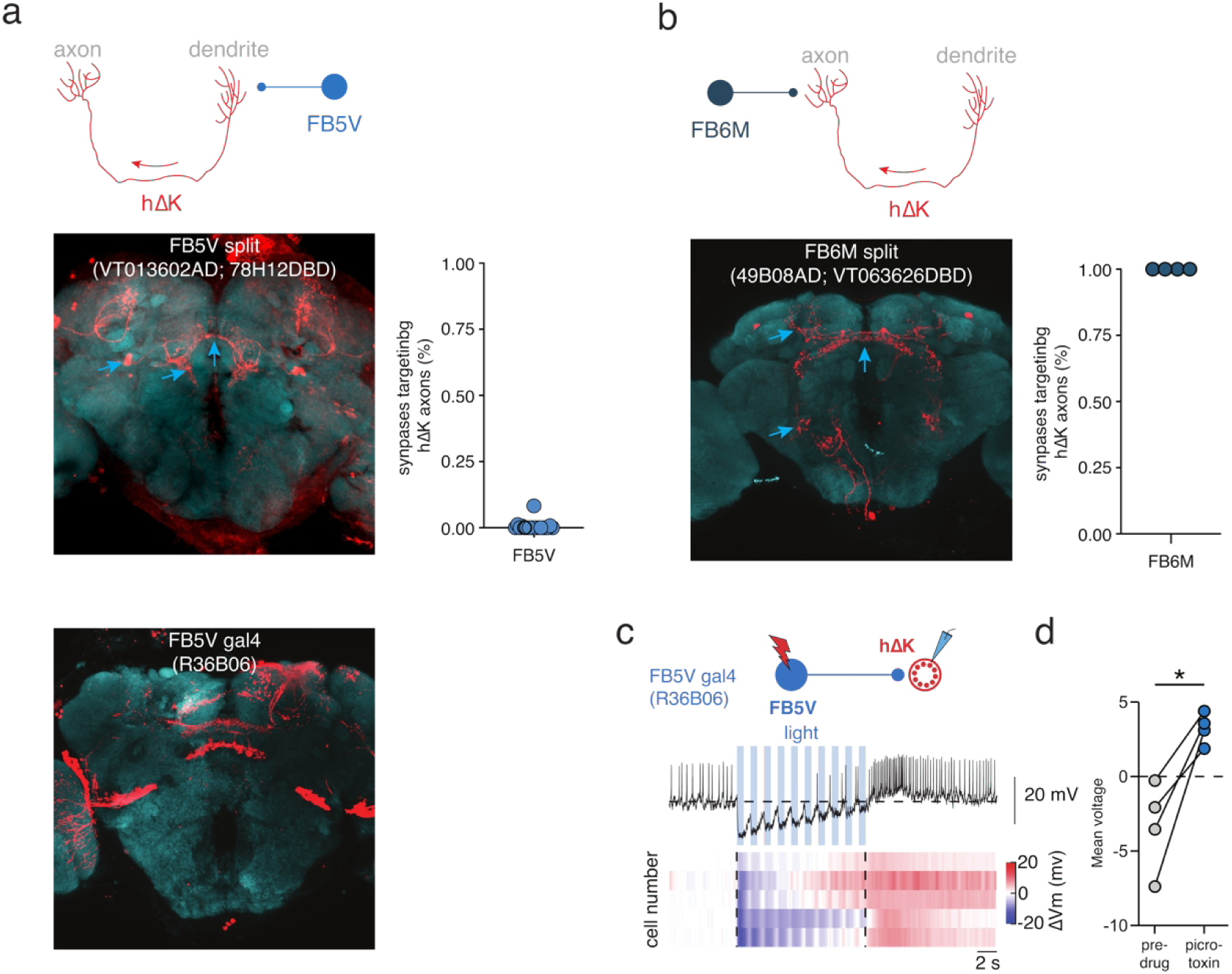
Additional data on inhibitory inputs to hΔK. **a,** Top, schematic showing axo-dendritic connectivity between FB5V and hΔK neurons. Middle, left, GFP staining of the line VT013602AD; 78H12DBD, which targets FB5V, among other neurons. Middle, right, percent of FB5V synapses targeting the hΔK axons. Each data point corresponds with one FB5V neuron. Bottom, GFP staining of the line R36B06-gal4, which targets FB5V, among other neurons. **b,** Top, schematic showing axo-axonic connectivity between FB6M and hΔK neurons. Bottom, left, GFP staining of the line 49B08AD; VT063626DBD, which targets FB6M. Arrows indicate characteristic morphological features of FB6M, including FB axons, ventral WED dendrites, and dorsal SMP dendrites. Bottom, right, percent of FB6M synapses targeting hΔK axons. Each data point corresponds with one FB6M neuron. **c**, Spiking and membrane potential response of hΔK neurons to optogenetic activation of FB5V (36B06-Gal4). Top: example trace; bottom: heat map of the mean membrane potential response with each cell as a row. **d**, Mean voltage responses of hΔK neurons in response to optogenetic activation of FB5V (36B06-Gal4) before and after the application of picrotoxin (5 µM).

**Ext. Data Fig. 5:**
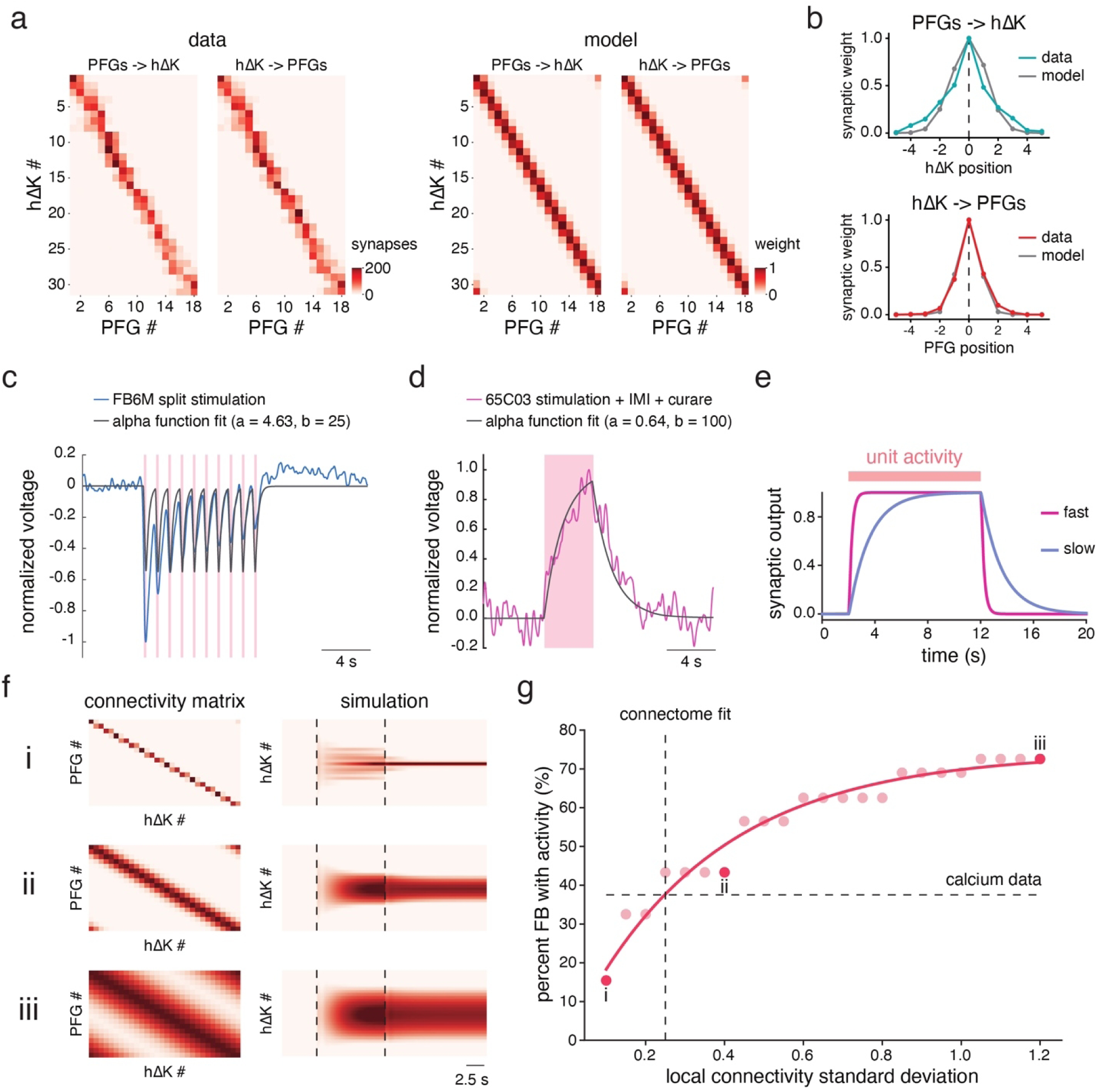
Methods details for recurrent model of hΔK and PFGs. **a,** Comparison of the real connectivity between hΔK and PFG neurons (left) and the Gaussian connectivity in the best fit model (right, σ = 0.25 radians). **b,** Local connectivity between hΔK and PFG neurons in the data and model, visualized by aligning each presynaptic neuron and averaging the synaptic weights to neighbouring postsynaptic partners. **c**, Alpha function fit to hΔK voltage responses to pulsed (0.2 duty cycle) stimulation of FB6M (49B08AD; VT063626DBD). **d**, Alpha function fit to hΔK voltage responses to a single 4 s stimulation of FB6A/D (65C03-gal4) in the presence of the nicotinic acetylcholine receptor blockers curare (10 µM) and IMI (100 nM). These blockers are included to remove acetylcholine signaling by FB6A/D and hΔK neurons, leaving only the slow synaptic signaling from FB6A/D for fitting. **e,** Synaptic output of model neurons using fast (pink) or slow (blue) synaptic speeds in response to a prolonged step of activity. **f**, Simulations of the hΔK-PFG network performed across various local connectivity widths. The Gaussian connectivity between hΔK and PFG neurons is shown on the left and the simulation results are shown on the right. Note how the bump width increases as the standard deviation of connectivity is increased. **g**, Width of attractor bumps (percent of the FB with activity) as a function of the standard deviation of recurrent connectivity (see Methods). Red line is an exponential plateau function fit to the simulation data (y = a*(1 – e^-bx^), a = 74, b = 2.83). The percent of FB with calcium activity during hΔK bumps and the local connectivity of the best fit Gaussian model are shown as dashed lines.

**Ext. Data Fig. 6:**
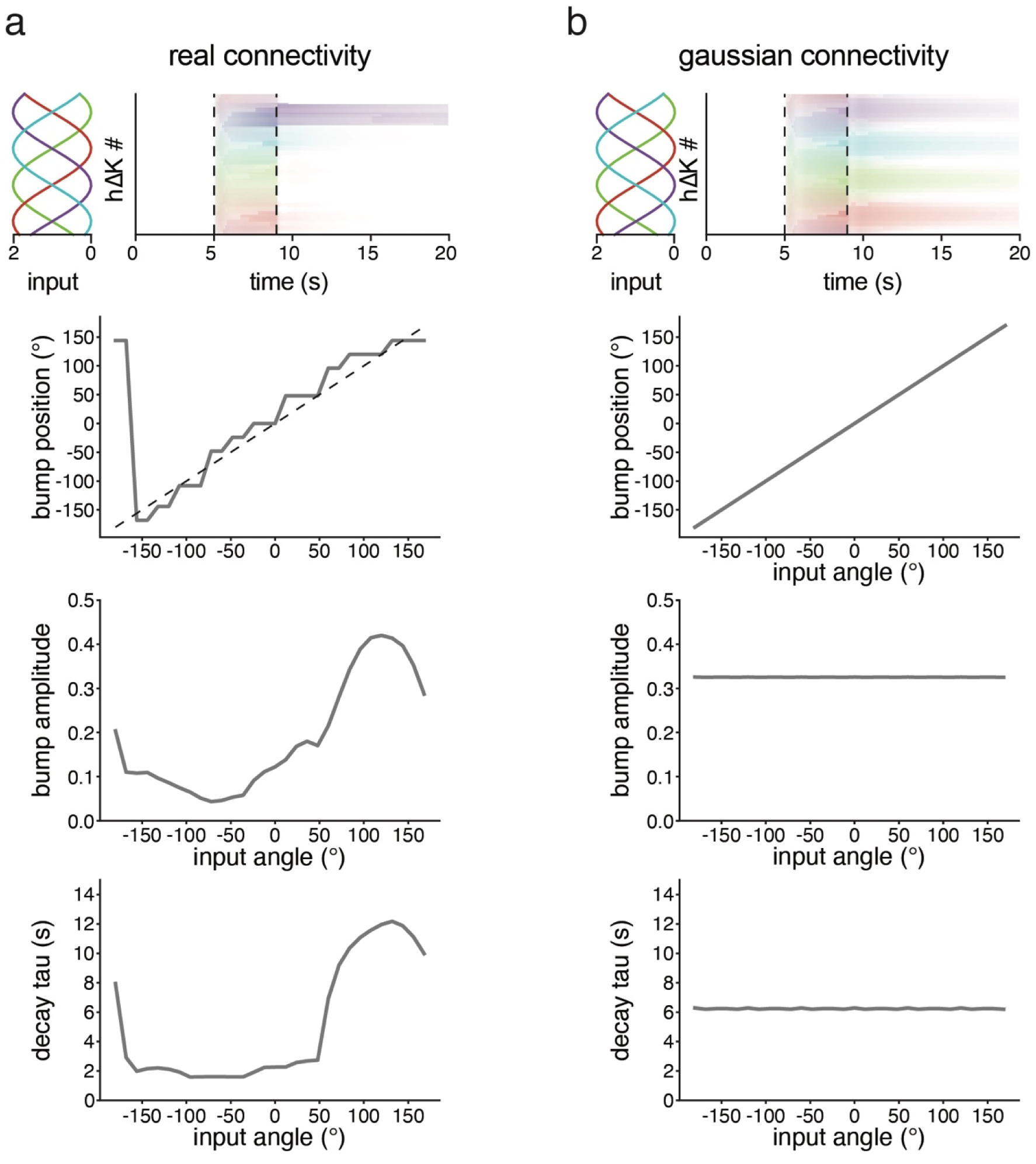
A model with real connectivity between PFG and hΔK can form a bump at most locations but shows inhomogeneity in bump amplitude and duration. **a,** hΔK bump position, bump amplitude, and decay rate (tau) in response to sinusoidal input across all possible phases when the connectivity between hΔK and PFG neurons is taken from the hemibrain connectome (see Ext. Data Fig. 1f). Top heatmap shows four separate simulations, all overlayed, in response to the sinusoidal inputs shown. Note that bump position can be stabilized at most positions, but the bump amplitude and decay time constant depend strongly on position. **b,** Same as **a** but using connections approximated by a Gaussian model.

**Ext. Data Fig. 7:**
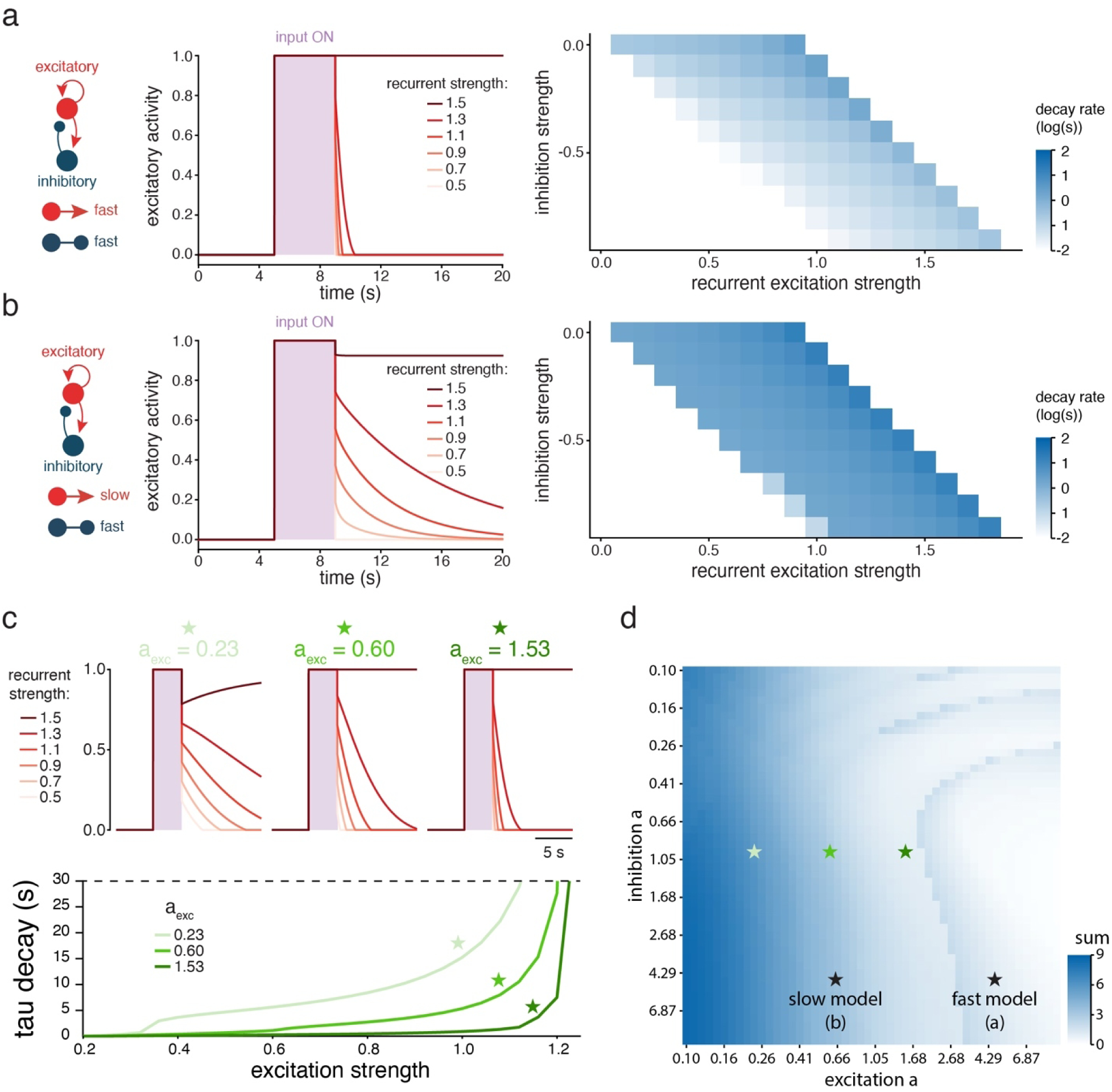
Slow excitation stabilizes persistent activity in a minimal recurrent network. **a,** Left, schematic of a simple two-unit recurrent network using fast excitation and slow excitation. Middle, activity of excitatory unit in response to inputs as a function of recurrent excitation strength. Right, parameter space showing the decay rate (log_10_ scale) of the excitatory unit following input offset as a function of recurrent excitation and inhibition strength. **b,** Same as **a** but the excitatory unit uses slow synaptic signaling. **c**, Two-unit model simulations as the excitatory synaptic decay rate (a) is slowed (a = 0.23, 0.60, or 1.53) and the recurrent excitatory strength is increased. Top, simulation output of the excitatory unit across conditions. Bottom, excitatory decay tau for each synaptic speed across recurrent excitation strengths. As the excitatory synaptic decay rate slows, the transition between fast activity decay and zero activity decay occurs more gradually. **d**, Parameter space showing the sum of the excitatory tau decay across excitation strengths (sum of each curve in **c**) across various speeds of excitation and inhibition. The models used in **a, b,** and **c** are all shown as stars.

**Ext. Data Fig. 8:**
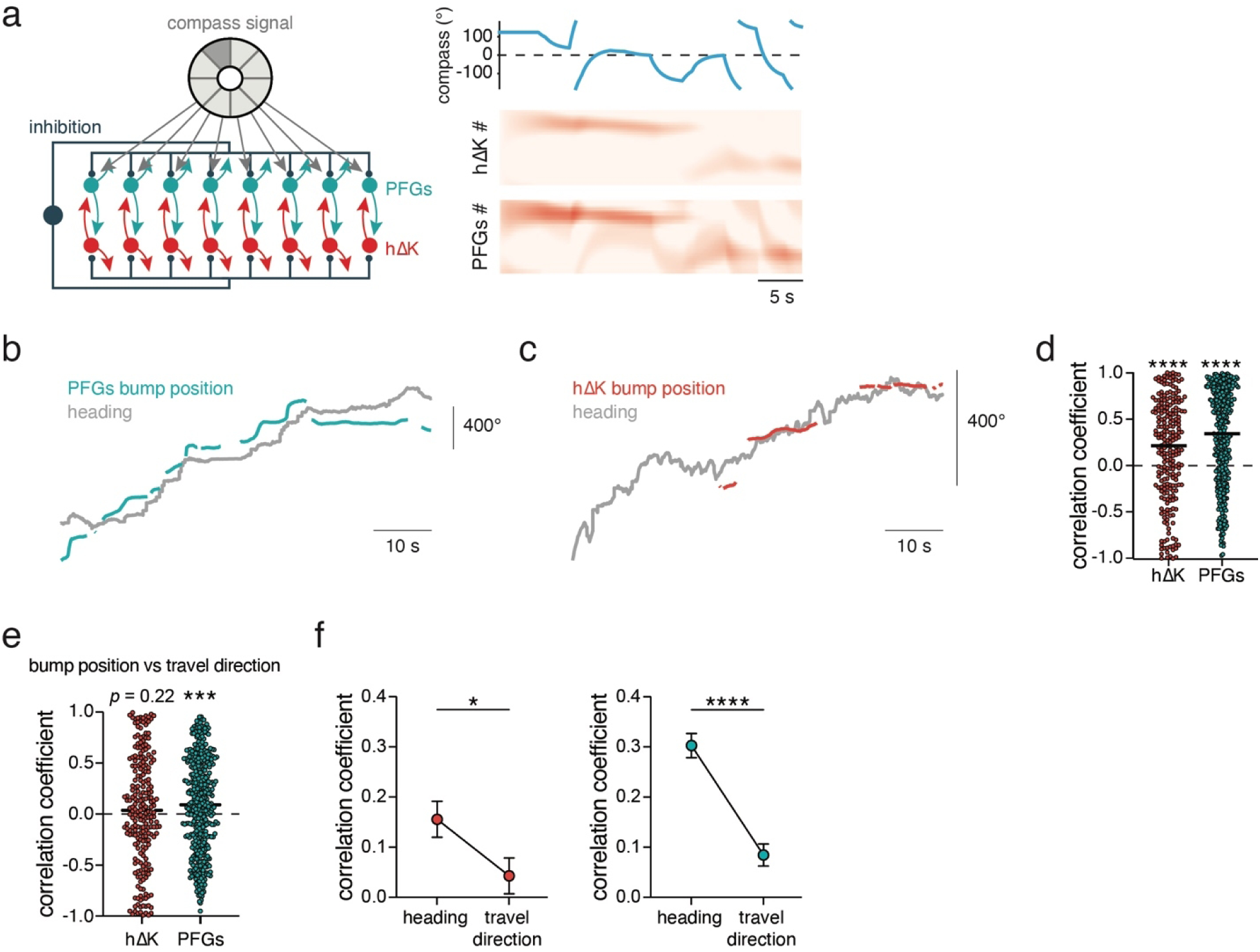
Additional data on differences in bump dynamics between PFG and hΔK. **a,** Simulations of hΔK and PFG population activity when the EPG compass signal (light blue) is randomly and stochastically updated and PFG neurons receive direct sinusoidal inputs from the compass. Left shows a schematic of the network configuration for the simulation. Right shows the simulation output. Note presence of a bump at all times in both populations, and occasionally the presence of multiple coexisting bumps in PFG neurons. **b,** Example of the PFG bump position aligned to the corresponding heading of the fly. **c,** Same as **b** but for hΔK neurons. **d,** Correlation coefficients between heading and hΔK (red) or PFG (blue) bump position across trials. **e,** Correlation coefficients between travel direction and hΔK (red) or PFG (blue) bump positions across trials. **f,** Average correlation coefficients of hΔK (left) or PFG (right) bump position with heading direction versus travel direction (error = mean ± SEM).

**Ext. Data Fig. 9:**
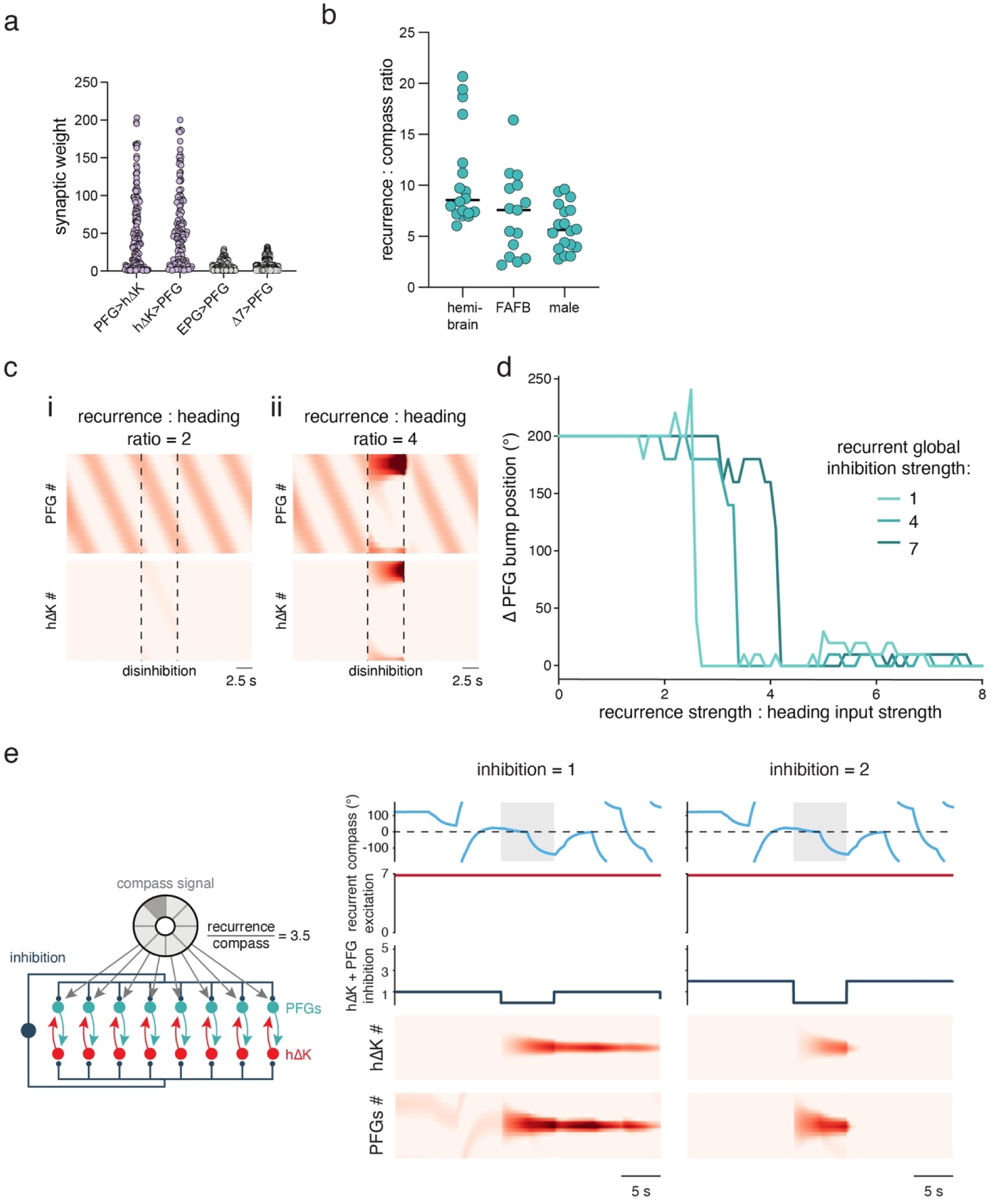
Additional data on disinhibition model. **a,** Synaptic weights between pairs of PFG and hΔK neurons, as well as from EPG and Δ7 neurons to PFG neurons. **b,** The ratio of the average synaptic input between hΔK neurons and PFG neurons (recurrent) to total synaptic input from EPGs (compass), for each PFG neuron. The average (mean ± SD) ratios for each connectome are 10.70 ± 4.816 (hemibrain), 7.15 ± 4.06 (FAFB), and 5.84 ± 2.23 (male CNS). **c**, Simulation of hΔK and PFG population activity with a disinhibitory gate of hΔK neurons when the recurrence : heading ratio is low (ratio = 2) or high (ratio = 7). The global recurrent inhibition strength is 1 for both simulations. Note the bump stabilization only occurs at the higher ratio. **d**, Change in PFG bump position during disinhibition as the recurrence : heading ratio increases, and as the recurrent global inhibition strength changes. Note the same heading signal is used during each simulation, which is a constant rotation, as seen in **c**. **e,** Simulation of hΔK and PFG population activity with inhibition to both PFG and hΔK. As in Fig. 5, EPG compass signal (light blue) is randomly and stochastically updated and global inhibition to hΔK and PFG neurons is transiently shut off. The global inhibition strength was set to 1 (left) or 2 (right), showing how the simulations gate activity in both hΔK and PFG neurons but do not permit continuous compass representations in PFG neurons during inhibition.

**Ext. Data Fig. 10:**
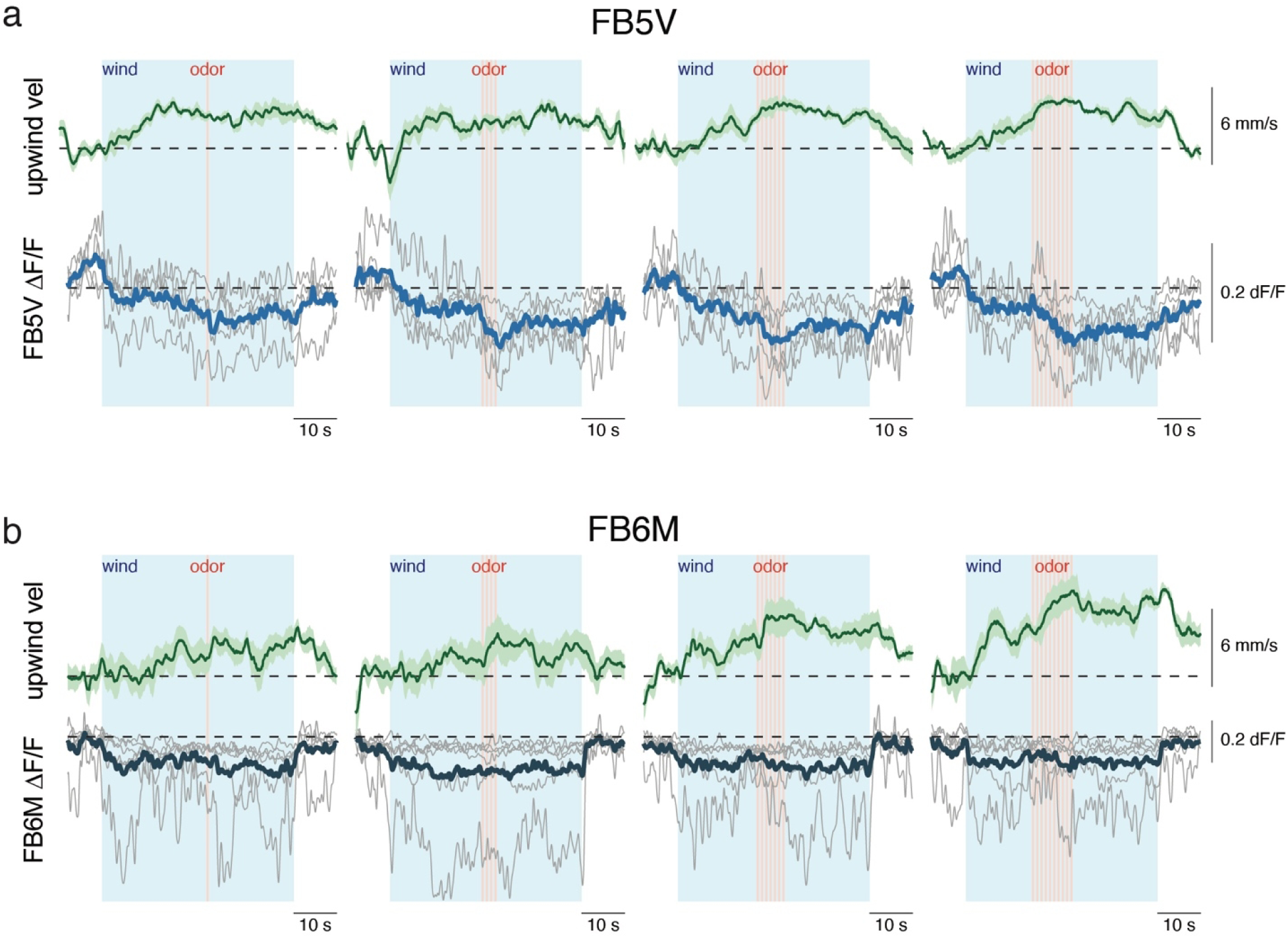
Additional data on encoding of sensory input by FB5V and FB6M. **a,** Mean upwind velocity and FB5V ΔF/F during baseline (no stimuli), wind, and pulses (1, 4, 7, or 10) of odor. Individual fly averages in grey; average across flies in green (behavior) and blue (neural activity). Shaded regions are mean ± SEM. **b,** Same as **a** but for FB6M neurons. Individual fly averages in grey; average across flies in green (behavior) and blue (neural activity).

## References

1. Compte, A., Brunel, N., Goldman-Rakic, P. S., & Wang, X. J. (2000). Synaptic mechanisms and network dynamics underlying spatial working memory in a cortical network model. Cerebral Cortex, 10(9), 910–923. 10.1093/cercor/10.9.910

2. Khona, M., & Fiete, I. R. (2022). Attractor and integrator networks in the brain. Nature Reviews Neuroscience, 23(12), 744–766. 10.1038/s41583-022-00642-0

3. Inagaki, H. K., Chen, S., Daie, K., Finkelstein, A., Fontolan, L., Romani, S., & Svoboda, K. (2022). Neural Algorithms and Circuits for Motor Planning. Annual Review of Neuroscience, 45, 249–271. 10.1146/annurev-neuro-092021-121730

4. Lo, C.-C., Boucher, L., Paré, M., Schall, J. D., & Wang, X.-J. (2009). Proactive Inhibitory Control and Attractor Dynamics in Countermanding Action: A Spiking Neural Circuit Model. Journal of Neuroscience, 29(28), 9059–9071. 10.1038/nn1722

5. Pereira, J., & Wang, X.-J. (2015). A Tradeoff Between Accuracy and Flexibility in a Working Memory Circuit Endowed with Slow Feedback Mechanisms. *Cerebral Cortex (New York*, N.Y.: 1991), *25*(10), 3586–3601. 10.1093/cercor/bhu202

6. Heeger, D. J., & Mackey, W. E. (2019). Oscillatory recurrent gated neural integrator circuits (ORGaNICs), a unifying theoretical framework for neural dynamics. Proceedings of the National Academy of Sciences of the United States of America, 116(45), 22783–22794. 10.1073/pnas.1911633116

7. Costa, R., Assael, I. A., Shillingford, B., de Freitas, N., & Vogels, T. (2017). Cortical microcircuits as gated-recurrent neural networks. Advances in Neural Information Processing Systems, 30. https://papers.nips.cc/paper_files/paper/2017/hash/45fbc6d3e05ebd93369ce542e8f2322d-Abstract.html

8. Hulse, B. K., Haberkern, H., Franconville, R., Turner-Evans, D., Takemura, S.-Y., Wolff, T., Noorman, M., Dreher, M., Dan, C., Parekh, R., Hermundstad, A. M., Rubin, G. M., & Jayaraman, V. (2021). A connectome of the Drosophila central complex reveals network motifs suitable for flexible navigation and context-dependent action selection—PubMed. eLife, 10, e66039.

9. Funahashi, S., Bruce, C. J., & Goldman-Rakic, P. S. (1989). Mnemonic coding of visual space in the monkey’s dorsolateral prefrontal cortex. Journal of Neurophysiology, 61(2), 331–349. 10.1152/jn.1989.61.2.331

10. Inagaki, H. K., Fontolan, L., Romani, S., & Svoboda, K. (2019). Discrete attractor dynamics underlies persistent activity in the frontal cortex. Nature, 566(7743), 212–217. 10.1038/s41586-019-0919-7

11. Claudi, F., Chandra, S., & Fiete, I. R. (2025). A theory and recipe to construct general and biologically plausible integrating continuous attractor neural networks. eLife, 14. 10.7554/eLife.107224.1

12. Wang, X.-J., & Yang, G. R. (2018). A disinhibitory circuit motif and flexible information routing in the brain. Current Opinion in Neurobiology, 49, 75–83. 10.1016/j.conb.2018.01.002

13. Kim, S. S., Rouault, H., Druckmann, S., & Jayaraman, V. (2017). Ring attractor dynamics in the Drosophila central brain. Science, 356(6340), 849–853. 10.1126/science.aal4835

14. Pang, M. M., Chen, F., Xie, M., Druckmann, S., Clandinin, T. R., & Yang, H. H. (2025). A recurrent neural circuit in Drosophila temporally sharpens visual inputs. Current Biology, 35(2), 333–346.e6. 10.1016/j.cub.2024.11.064

15. Christenson, M. P., Sanz Diez, A., Heath, S. L., Saavedra-Weisenhaus, M., Adachi, A., Nern, A., Abbott, L. F., & Behnia, R. (2024). Hue selectivity from recurrent circuitry in Drosophila. Nature Neuroscience, 27(6), 1137–1147. 10.1038/s41593-024-01640-4

16. Scheffer, L. K., Xu, C. S., Januszewski, M., Lu, Z., Takemura, S.-Y., Hayworth, K. J., Huang, G. B., Shinomiya, K., Maitlin-Shepard, J., Berg, S., Clements, J., Hubbard, P. M., Katz, W. T., Umayam, L., Zhao, T., Ackerman, D., Blakely, T., Bogovic, J., Dolafi, T., … Plaza, S. M. (2020). A connectome and analysis of the adult Drosophila central brain. eLife, 9, e57443. 10.7554/eLife.57443

17. Dorkenwald, S., Matsliah, A., Sterling, A. R., Schlegel, P., Yu, S.-C., McKellar, C. E., Lin, A., Costa, M., Eichler, K., Yin, Y., Silversmith, W., Schneider-Mizell, C., Jordan, C. S., Brittain, D., Halageri, A., Kuehner, K., Ogedengbe, O., Morey, R., Gager, J., … FlyWire Consortium. (2024). Neuronal wiring diagram of an adult brain. Nature, 634(8032), 124–138. 10.1038/s41586-024-07558-y

18. Wolff, T., Eddison, M., Chen, N., Nern, A., Sundaramurthi, P., Sitaraman, D., & Rubin, G. M. (2025). Cell type-specific driver lines targeting the Drosophila central complex and their use to investigate neuropeptide expression and sleep regulation. eLife, 14, RP104764. 10.7554/eLife.104764

19. Stone, T., Webb, B., Adden, A., Weddig, N. B., Honkanen, A., Templin, R., Wcislo, W., Scimeca, L., Warrant, E., & Heinze, S. (2017). An Anatomically Constrained Model for Path Integration in the Bee Brain. Current Biology: CB, 27(20), 3069–3085.e11. 10.1016/j.cub.2017.08.052

20. Lu, J., Behbahani, A. H., Hamburg, L., Westeinde, E. A., Dawson, P. M., Lyu, C., Maimon, G., Dickinson, M. H., Druckmann, S., & Wilson, R. I. (2022). Transforming representations of movement from body- to world-centric space. Nature, 601(7891), 98–104. 10.1038/s41586-021-04191-x

21. Lyu, C., Abbott, L. F., & Maimon, G. (2022). Building an allocentric travelling direction signal via vector computation. Nature, 601(7891), 92–97. 10.1038/s41586-021-04067-0

22. Matheson, A. M. M., Lanz, A. J., Medina, A. M., Licata, A. M., Currier, T. A., Syed, M. H., & Nagel, K. I. (2022). A neural circuit for wind-guided olfactory navigation. Nature Communications, 13(1), 4613. 10.1038/s41467-022-32247-7

23. Beetz, M. J., Kraus, C., & El Jundi, B. (2023). Neural representation of goal direction in the monarch butterfly brain. Nature Communications, 14(1), 5859. 10.1038/s41467-023-41526-w

24. Mussells Pires, P., Zhang, L., Parache, V., Abbott, L. F., & Maimon, G. (2024). Converting an allocentric goal into an egocentric steering signal. Nature, 626(8000), 808–818. 10.1038/s41586-023-07006-3

25. Westeinde, E. A., Kellogg, E., Dawson, P. M., Lu, J., Hamburg, L., Midler, B., Druckmann, S., & Wilson, R. I. (2024). Transforming a head direction signal into a goal-oriented steering command. Nature, 626(8000), 819–826. 10.1038/s41586-024-07039-2

26. Kathman, N. D., Lanz, A. J., Freed, J. D., & Nagel, K. I. (2025). Neural dynamics for working memory and evidence integration during olfactory navigation in Drosophila (p. 2024.10.05.616803). bioRxiv. 10.1101/2024.10.05.616803

27. Siliciano, A. F., Minni, S., Morton, C., Dowell, C. K., Eghbali, N. B., Rhee, J. Y., Abbott, L. F., & Ruta, V. (2025). A vector-based strategy for olfactory navigation in Drosophila. 10.1101/2025.02.15.638426

28. Flores-Valle, A., Honnef, R., & Seelig, J. D. (2025). Goal learning, memory, and drift in the Drosophila head direction system (p. 2025.03.20.644317). bioRxiv. 10.1101/2025.03.20.644317

29. Seelig, J. D., & Jayaraman, V. (2015). Neural dynamics for landmark orientation and angular path integration. Nature, 521(7551), 186–191. 10.1038/nature14446

30. Demir, M., Kadakia, N., Anderson, H. D., Clark, D. A., & Emonet, T. (2020). Walking Drosophila navigate complex plumes using stochastic decisions biased by the timing of odor encounters. eLife, 9, e57524. 10.7554/eLife.57524

31. Eckstein, N., Bates, A. S., Champion, A., Du, M., Yin, Y., Schlegel, P., Lu, A. K.-Y., Rymer, T., Finley-May, S., Paterson, T., Parekh, R., Dorkenwald, S., Matsliah, A., Yu, S.-C., McKellar, C., Sterling, A., Eichler, K., Costa, M., Seung, S., … Funke, J. (2024). Neurotransmitter classification from electron microscopy images at synaptic sites in Drosophila melanogaster. Cell, 187(10), 2574–2594.e23. 10.1016/j.cell.2024.03.016

32. Nagel, K. I., Hong, E. J., & Wilson, R. I. (2015). Synaptic and circuit mechanisms promoting broadband transmission of olfactory stimulus dynamics. Nature Neuroscience, 18(1), 56–65. 10.1038/nn.3895

33. Liu, W. W., Wilson, R. I. (2013). Glutamate is an inhibitory neurotransmitter in the Drosophila olfactory system. Proceedings of the National Academy of Sciences of the United States of America, 110(25), 10294–10299. 10.1073/pnas.1220560110

34. Omoto, J. J., Nguyen, B. C. M., Kandimalla, P., Lovick, J. K., Donlea, J. M., & Hartenstein, V. (2018). Neuronal constituents and putative interactions within the Drosophila ellipsoid body neuropil. Frontiers in neural circuits, 12, 103.

35. Wang, X. J. (1999). Synaptic basis of cortical persistent activity: The importance of NMDA receptors to working memory. The Journal of Neuroscience: The Official Journal of the Society for Neuroscience, 19(21), 9587–9603. 10.1523/JNEUROSCI.19-21-09587.1999

36. Noorman, M., Hulse, B. K., Jayaraman, V., Romani, S., & Hermundstad, A. M. (2024). Maintaining and updating accurate internal representations of continuous variables with a handful of neurons. Nature Neuroscience, 27(11), 2207–2217.

37. Green, J., Adachi, A, Shah, K. K., Hirokawa, J. D., Magani, P. S., & Maimon, G. (2017). A neural circuit architecture for angular integration in Drosophila. Nature, 546(7656), 101–106. doi: 10.1038/nature22343

38. Turner-Evans, D., Wegener, S., Rouault, H., Franconville, R., Wolff, T., Seelig, J. D., Druckmann, Shaul, & Jayaraman, V. (2017). Angular velocity integration in a fly heading circuit. Elife, 6, e23496. doi: 10.7554/eLife.23496

39. Scaplen, K. M., Talay, M., Fisher, J. D., Cohn, R., Sorkaç, A., Aso, Y., Barnea, G., & Kaun, K. R. (2021). Transsynaptic mapping of Drosophila mushroom body output neurons. eLife, 10, e63379. 10.7554/eLife.63379

40. Namiki, S., & Kanzaki, R. (2016). Comparative neuroanatomy of the lateral accessory lobe in the insect brain. Frontiers in physiology, 7, 244.

41. Rayshubskiy, A., Holtz, S. L., Bates, A. S., Vanderbeck, Q. X., Capdevila, L. S., Rockwell, V., & Wilson, R. (2025). Neural circuit mechanisms for steering control in walking Drosophila. ELife, 13, RP102230.

42. Feng, K., Khan, M., Minegishi, R., Müller, A., Poll, M. N. V. D., Swinderen, B. van, & Dickson, B. J. (2024). A central steering circuit in Drosophila (p. 2024.06.27.601106). bioRxiv. 10.1101/2024.06.27.601106

43. Patella, P., & Wilson, R. I. (2018). Functional Maps of Mechanosensory Features in the Drosophila Brain. Current Biology, 28(8), 1189–1203.e5. 10.1016/j.cub.2018.02.074

44. Suver, M. P., Matheson, A. M. M., Sarkar, S., Damiata, M., Schoppik, D., & Nagel, K. I. (2019). Encoding of Wind Direction by Central Neurons in Drosophila. Neuron, 102(4), 828–842.e7. 10.1016/j.neuron.2019.03.012

45. Seung, H. S. (1996). How the brain keeps the eyes still. Proceedings of the National Academy of Sciences of the United States of America, 93(23), 13339–13344. 10.1073/pnas.93.23.13339

46. Vishwanathan, A., Sood, A., Wu, J., Ramirez, A. D., Yang, R., Kemnitz, N., Ih, D., Turner, N., Lee, K., Tartavull, I., Silversmith, W. M., Jordan, C. S., David, C., Bland, D., Sterling, A., Seung, H. S., Goldman, M. S., Aksay, E. R. F., & Eyewirers. (2024). Predicting modular functions and neural coding of behavior from a synaptic wiring diagram. Nature Neuroscience, 27(12), 2443–2454. 10.1038/s41593-024-01784-3

47. Burak, Y., & Fiete, I. R. (2009). Accurate path integration in continuous attractor network models of grid cells. PLoS Computational Biology, 5(2), e1000291. 10.1371/journal.pcbi.1000291

48. Kennedy, A., Kunwar, P. S., Li, L.-Y., Stagkourakis, S., Wagenaar, D. A., & Anderson, D. J. (2020). Stimulus-specific hypothalamic encoding of a persistent defensive state. Nature, 586(7831), 730–734. 10.1038/s41586-020-2728-4

49. Liu, M., Nair, A., Coria, N., Linderman, S. W., & Anderson, D. J. (2024). Encoding of female mating dynamics by a hypothalamic line attractor. Nature, 634(8035), 901–909. 10.1038/s41586-024-07916-w

50. Litwin-Kumar, A., & Doiron, B. (2014). Formation and maintenance of neuronal assemblies through synaptic plasticity. Nature Communications, 5, 5319. 10.1038/ncomms6319

51. Kutschireiter, A., Basnak, M. A., Wilson, R. I., & Drugowitsch, J. (2023). Bayesian inference in ring attractor networks. Proceedings of the National Academy of Sciences of the United States of America, 120(9), e2210622120. 10.1073/pnas.2210622120

52. Daie, K., Fontolan, L., Druckmann, S., & Svoboda, K. (2023). Feedforward amplification in recurrent networks underlies paradoxical neural coding (p. 2023.08.04.552026). bioRxiv. 10.1101/2023.08.04.552026

53. Masse, N. Y., Yang, G. R., Song, H. F., Wang, X.-J., & Freedman, D. J. (2019). Circuit mechanisms for the maintenance and manipulation of information in working memory. Nature Neuroscience, 22(7), 1159–1167. 10.1038/s41593-019-0414-3

54. Pi, H.-J., Hangya, B., Kvitsiani, D., Sanders, J. I., Huang, Z. J., & Kepecs, A. (2013). Cortical interneurons that specialize in disinhibitory control. Nature, 503(7477), 521–524. 10.1038/nature12676

55. Keller, A. J., Dipoppa, M., Roth, M. M., Caudill, M. S., Ingrosso, A., Miller, K. D., & Scanziani, M. (2020). A Disinhibitory Circuit for Contextual Modulation in Primary Visual Cortex. Neuron, 108(6), 1181–1193.e8. 10.1016/j.neuron.2020.11.013

56. Krabbe, S., Paradiso, E., d’Aquin, S., Bitterman, Y., Courtin, J., Xu, C., Yonehara, K., Markovic, M., Müller, C., Eichlisberger, T., Gründemann, J., Ferraguti, F., & Lüthi, A. (2019). Adaptive disinhibitory gating by VIP interneurons permits associative learning. Nature Neuroscience, 22(11), 1834–1843. 10.1038/s41593-019-0508-y

57. Hikosaka, O., & Wurtz, R. H. (1983). Visual and oculomotor functions of monkey substantia nigra pars reticulata. IV. Relation of substantia nigra to superior colliculus. Journal of Neurophysiology, 49(5), 1285–1301. 10.1152/jn.1983.49.5.1285

58. Falasconi, A., Kanodia, H., & Arber, S. (2025). Dynamic basal ganglia output signals license and suppress forelimb movements. Nature. 10.1038/s41586-025-09066-z

59. Manu, M., & Baccus, S. A. (2011). Disinhibitory gating of retinal output by transmission from an amacrine cell. Proceedings of the National Academy of Sciences of the United States of America, 108(45), 18447–18452. 10.1073/pnas.1107994108

60. Gale, S. D., & Perkel, D. J. (2010). A basal ganglia pathway drives selective auditory responses in songbird dopaminergic neurons via disinhibition. The Journal of Neuroscience, 30(3), 1027–1037. 10.1523/JNEUROSCI.3585-09.2010

61. Kosche, G., Vallentin, D., & Long, M. A. (2015). Interplay of inhibition and excitation shapes a premotor neural sequence. The Journal of Neuroscience: The Official Journal of the Society for Neuroscience, 35(3), 1217–1227. 10.1523/JNEUROSCI.4346-14.2015

62. Wen, X., Connors, K. E., Snell, C. C., McGrath, R., Mehta, A., & Hattori, D. (2004). A dopamine circuit regulates locomotor initiation and persistence in *Drosophil* (p. 2025.11.18.689052). *bioRxiv*. doi: 10.1101/2025.11.18.689052

63. Constantinidis, C., & Wang, X.-J. (2004). A neural circuit basis for spatial working memory. The Neuroscientist, 10(6), 553–565. 10.1177/1073858404268742

64. Vinograd, A., Nair, A., Kim, J. H., Linderman, S. W., & Anderson, D. J. (2024). Causal evidence of a line attractor encoding an affective state. Nature, 634(8035), 910–918. 10.1038/s41586-024-07915-x

65. Mountoufaris, G., Nair, A., Yang, B., Kim, D.-W., Vinograd, A., Kim, S., Linderman, S. W., & Anderson, D. J. (2024). A line attractor encoding a persistent internal state requires neuropeptide signaling. Cell, 187(21), 5998–6015.e18. 10.1016/j.cell.2024.08.015

66. Epiney, D. G., Chaya, G. M., Dillon, N. R., Lai, S.-L., & Doe, C. Q. (2025). Single nuclei RNA-sequencing of adult brain neurons derived from type 2 neuroblasts reveals transcriptional complexity in the insect central complex. eLife, 14, RP105896. 10.7554/eLife.105896

67. Kleinfeld, D., Raccuia-Behling, F., & Chiel, H. J. (1990). Circuits constructed from identified Aplysia neurons exhibit multiple patterns of persistent activity. Biophysical journal, 57(4), 697–715.

68. Fisher, Y. E., Marquis, M., D’Alessandro, I., & Wilson, R. I. (2022). Dopamine promotes head direction plasticity during orienting movements. Nature, 612(7939), 316–322. 10.1038/s41586-022-05485-4

69. Le Moël, F., Stone, T., Lihoreau, M., Wystrach, A., & Webb, B. (2019). The Central Complex as a Potential Substrate for Vector Based Navigation. Frontiers in Psychology, 10, 690. 10.3389/fpsyg.2019.00690

70. Dan, C., Hulse, B. K., Kappagantula, R., Jayaraman, V., & Hermundstad, A. M. (2024). A neural circuit architecture for rapid learning in goal-directed navigation. Neuron, 112(15), 2581–2599.e23. 10.1016/j.neuron.2024.04.036

71. Cho, K., van Merriënboer, B., Bahdanau, D., & Bengio, Y. (2014). On the Properties of Neural Machine Translation: Encoder–Decoder Approaches. In D. Wu, M. Carpuat, X. Carreras, & E. M. Vecchi (Eds.), Proceedings of SSST-8, Eighth Workshop on Syntax, Semantics and Structure in Statistical Translation (pp. 103–111). Association for Computational Linguistics. 10.3115/v1/W14-4012

72. Krishnamurthy, K., Can, T., & Schwab, D. J.. (2022). Theory of Gating in Recurrent Neural Networks. Physical Review X, 12. https://journals.aps.org/prx/abstract/10.1103/PhysRevX.12.011011

73. Driscoll, L. N., Shenoy, K., & Sussillo, D. (2024). Flexible multitask computation in recurrent networks utilizes shared dynamical motifs. Nature Neuroscience, 27(7), 1349–1363. 10.1038/s41593-024-01668-6

74. Sen, R., Wang, K., & Dickson, B. J. (2019). TwoLumps Ascending Neurons Mediate Touch-Evoked Reversal of Walking Direction in Drosophila. Current Biology, 29(24), 4337–4344. doi: 10.1016/j.cub.2019.11.004

75. Sherer, L. M., Garrett, E. C., Morgan, H. R., Brewer, E. D., Sirrs, L. A., Shearin, H. K., Williams, J. L., McCabe, B. D., Stowers, R. S., & Cartel, S. J. (2020). Octopamine neuron dependent aggression requires dVGLUT from dual-transmitting neurons. PLos Genetics, 16(2), e1008609. doi: 10.1371/journal.pgen.1008609.

76. Hamid, A., Gattuso, H. Caglar, A. N., Pillai, M., Steele, T., Gonzalez, A., Nagel, K., & Syed, M. H. (2024). The conserved RNA-binding protein Imp is required for the specification and function of olfactory navigation circuitry in Drosophila. Current Biology, 34(3), 473–488. doi: 10.1016/j.cub.2023.12.020

77. Moore, R. J. D., Taylor, G. J., Paulk, A. C., Pearson, T., van Swinderen, B., & Srinivasan M. V. (2014). FicTrac: A visual method for tracking spherical motion and generating fictive animal paths. J Neurosci Methods, 225: 106–119. doi: 10.1016/j.jneumeth.2014.01.010 J Neurosci Methods225

78. Jenett, A., Rubin, G. M., Ngo, T.-T. B., Shepherd, D., Murphy, C., Dionne, H., Pfeiffer, B. D., Cavallaro, A., Hall, D., Jeter, J., Iyer, N., Fetter, D., Hausenfluck, J. H., Peng, H., Trautman, E. T., Svirskas, R. R., Myers, E. W., Iwinski, Z. R., Aso, Y., … Zugates, C. T. (2012). A GAL4-Driver Line Resource for *Drosophila* Neurobiology. Cell Reports, 2(4), 991–1001. 10.1016/j.celrep.2012.09.011

79. Bogovic, J. A., Otsuna, H., Heinrich, L., Ito, M., Jeter, J., Meissner, G., Nern, A., Colonell, J., Malkesman, O., Ito, K., & Saalfeld, S. (2020). An unbiased template of the Drosophila brain and ventral nerve cord. PLOS ONE, 15(12), e0236495. 10.1371/journal.pone.0236495

